# Single-cell transcriptome atlas of *Drosophila* gastrula 2.0

**DOI:** 10.1101/2021.12.27.474293

**Authors:** Shunta Sakaguchi, Yasushi Okochi, Chiharu Tanegashima, Osamu Nishimura, Tadashi Uemura, Mitsutaka Kadota, Honda Naoki, Takefumi Kondo

**Author notes:** To whom correspondence should be addressed. Takefumi Kondo, Ph.D.

## Abstract

During development, positional information directs cells to specific fates, leading them to differentiate with their own transcriptomes and express specific behaviors and functions. However, the mechanisms underlying these processes in a genome-wide view remain ambiguous, partly because the single-cell transcriptomic data of early developing embryos containing both accurate spatial and lineage information is still lacking. Here, we report a new single-cell transcriptome atlas of *Drosophila* gastrulae, divided into 65 transcriptomically distinct clusters. We found that the expression profiles of plasma-membrane-related genes, but not those of transcription factor genes, represented each germ layer, supporting the nonequivalent contribution of each transcription factor mRNA level to effector gene expression profiles at the transcriptome level. We also reconstructed the spatial expression patterns of all genes at the single-cell stripe level as the smallest unit. This atlas is an important resource for the genome-wide understanding of the mechanisms by which genes cooperatively orchestrate *Drosophila* gastrulation.

## Introduction

One of the fundamental goals of developmental biology is to understand how genes cooperatively orchestrate morphogenesis and physiological functions at the cellular and tissue levels. During embryogenesis, a fertilized egg divides multiple times to increase the cell number. The proliferating cells form blastoderm, and the embryo undergoes gastrulation and later organogenesis (Perrimon et al., 2012). In the scheme of programmed control of development, positional information is established by the combination of morphogens and cell-cell interactions, and each cell is canalized into a specific fate depending on their positions in embryos (Collinet and Lecuit, 2021). Initial positional information is often graded and continuous, but the consequence of cell fate determination shows a discrete spatial pattern (Briscoe and Small, 2015; Petkova et al., 2019). The status of cell fates is often recognized by the expression of a limited number of marker genes, most of which encode transcription factors (TFs). It is widely considered that the combinatorial action of TFs through the dynamics of gene regulatory networks is important for transforming analog gradual information into a discrete digital pattern of gene expression (Briscoe and Small, 2015; Small and Arnosti, 2020; Zinzen et al., 2009), and new mechanisms to achieve this non-linear transformation have also been recently reported (Lammers et al., 2020; Papadopoulos and Tomancak, 2019). After cell fate canalization, each cell behaves differently to drive multicellular tissue morphogenesis (Heisenberg and Bellaiche, 2013; Kondo and Hayashi, 2015). However, since most studies have only considered a limited number of marker genes and focused on specific target genes/enhancers, genome-wide insight into cell differentiation and morphogenesis is still limited. To overcome these limitations, it is important to establish a single-cell transcriptome atlas of early developing embryos that contains both accurate spatial and lineage information.

*Drosophila* gastrulae has been an excellent model system for studying multicellular morphogenesis for decades (Gheisari et al., 2020; Martin, 2020; Wieschaus and Nüsslein-Volhard, 2016). The fertilized egg undergoes 13 rounds of nuclear division and then forms a single layer of the epithelial sheet covering the entire embryo, the so-called blastoderm. Although each cell has a similar columnar shape at this time, positional information along the anterior-posterior axis and dorsal-ventral axis has already been established in embryos and directs cells to express different set cell fate determinants (Jaeger, 2011; Perrimon et al., 2012). The cell-specific expression of genes has been extensively analyzed using *in situ* hybridization (ISH), but it is difficult to obtain the transcriptome profiles of each cell with spatial information from these data. For example, more than ten thousand genes have been analyzed by fluorescent ISH (FISH), but they are far from quantification and cannot integrate all FISH data per cell (Lécuyer et al., 2007; Wilk et al., 2016). The Berkeley Drosophila Transcription Network Project (BDTNP) established a gene expression database as a virtual embryo by integrating the quantified FISH data from multiple embryos, but the number of genes analyzed was less than 100 (Fowlkes et al., 2008; Keränen et al., 2006; Luengo Hendriks et al., 2006).

In this decade, single-cell RNA-sequencing (scRNA-seq) has become a standard technique that enables the analysis of transcriptomes at the single-cell level (Kolodziejczyk et al., 2015; Tanay and Regev, 2017). One of the disadvantages of scRNA-seq for tissue or embryo scale analysis is that each data loses the information of the original position of each cell in tissue, because, in conventional scRNA-seq protocols, cells need to be dissociated into the single-cell level from tissues. Several computational methods have been proposed to restore the spatial information of scRNA-seq data (Achim et al., 2015; Satija et al., 2015; Stuart et al., 2019; Welch et al., 2019). Basically, these methods can infer the original position of every cell from each scRNA-seq data by statistically comparing it to the known spatial expression profile of some landmark genes and thereby reconstruct the spatial profiles of all genes based on the inferred position. For *Drosophila* gastrulae, scRNA-seq analysis and spatial reconstruction of gene expression were performed (Karaiskos et al., 2017). However, there is room for improvement in its quality because, for many genes, the reconstructed spatial pattern from scRNA-seq data did not match the original pattern uncovered by *in situ* hybridization. For example, pair-rule genes (e.g., fushi tarazu (*ftz*), even-skipped (*eve*)), and segment polarity genes (e.g., wingless (*wg*), engrailed (*en*)) show seven- or fourteen-stripe expression patterns along the anterior-posterior axis in the gastrulae, but 14 stripes of segment polarity genes have not been fully reconstructed in the *Drosophila* Virtual Expression eXplorer (DVEX; http://DVEX.org). Recently, we computationally overcome this limitation by developing a new machine learning method, Perler, based on generative linear mapping (Okochi et al., 2021). Using the same scRNA-seq dataset with DVEX, Perler can reconstruct 14 stripe patterns of segment polarity genes. Furthermore, Perler has a notable feature in that the scRNA-seq data are not overfitted to the reference FISH data.

Although we and others have made efforts to improve computational methods (Cang and Nie, 2020; Nitzan et al., 2019; Tanevski et al., 2020), there is a fundamental limitation with the scRNA-seq data itself in using their outputs for biological research; the number of high-quality cells sequenced was 1,297, which is much lower than the number of cells in a gastrula (approximately 6,000 cells) (Karaiskos et al., 2017). Even though the gastrula is bilaterally symmetric and can be thought of as half 3,000 cells, there might be cells not in the scRNA-seq data. Karaiskos et al. were able to distinguish only 13 clusters in their scRNA-seq data, which may not be sufficient to fully describe the entire *Drosophila* gastrula. In addition, the depth of sequenced reads per cell was limited. Deeper data will allow us to understand the characteristics of each cell in more detail. Therefore, new acquisition of scRNA-seq data with higher numbers or depth will enable us to perform a more precise transcriptomic characterization, and the improvement of the spatial transcriptome reconstruction, which are of critical importance for understanding the mechanisms of cell differentiation and tissue morphogenesis during gastrulation from a genome-wide perspective.

In this study, we aimed to improve the single-cell transcriptome atlas of *Drosophila* gastrula and update it into version 2.0. To this end, we first developed a scRNA-seq protocol for *Drosophila* embryos with single-cell dissociation using cold-active protease (CAP), followed by non-crosslinked fixation of cells. CAP dissociation allows us to maintain the original gene expression profiles better than canonical trypsin dissociation. Second, we profiled single-cell transcriptomes with a higher number of cells and greater sequence depths, and manually annotated them into 65 clusters. Using this dataset, we revealed that the expression profiles of plasma membrane and cytoplasmic protein genes represent the cellular characteristics, such as differences between the three germ layers, better than those of the nuclear transcription factor genes. By detailed clustering analysis using previous biological knowledge, we further recapitulated the transcriptome profiles of each pair-rule stripe. Finally, we cataloged the spatial expression patterns of all genes in *Drosophila* gastrulae via computational integration with reference spatial expression patterns using Perler (Okochi et al., 2021) or NovoSpaRc (Moriel et al., 2021; Nitzan et al., 2019). Since *Drosophila* gastrula is one of the most well-characterized multicellular systems, this new atlas provides an important quantitative resource for a wide range of biological fields as a reference for understanding the principles that link gene regulatory networks and cell differentiation to cell behavior and tissue morphogenesis.

## Results

### Single-cell RNA-seq of fixed-cells dissociated from *Drosophila* embryos

To conduct scRNA-seq analysis for *Drosophila* gastrulae, we first reexamined the commonly used protocols of the single-cell dissociation step. We tried the mechanical dissociation protocol as previously reported (Karaiskos et al., 2017), but did not recover enough cells in our hands. Therefore, we next attempted gentle breaking of the vitelline membrane and dissociated the cells enzymatically. It has recently been recognized that enzymatic dissociation at room temperature leads to artificial changes in the transcriptome. To overcome this problem, the use of CAP has recently been shown to be a good solution (Adam et al., 2017; Denisenko et al., 2020; Miyawaki-Kuwakado et al., 2021; O’Flanagan et al., 2019). CAP (also known as subtilisin A from *Bacillus licheniformis*) is active even at 6 °C and can dissociate cells with minimal impact on gene expression. To assess the artificial effect of trypsin and the usefulness of CAP on single-cell dissociation of *Drosophila* embryos, we examined both enzymes (Figure 1A). In addition, in the step of isolating single cells using microfluidic devices, such as Fluidigm C1 (C1) or 10x Chromium, it is assumed that the live cells are exposed to room temperature, and this procedure may compromise the gene expression of the cells. Furthermore, in the case of C1, by inspecting cells after loading and capturing, we noticed that cells often undergo cell death in the C1 chip after loading, possibly due to mechanical stresses caused by flowing through microfluidic channels. To solve these problems, we added a step of non-cross-linking fixation using CellCover after dissociation (Figure 1A). To evaluate the preservation efficiency of fixation, we performed bulk RNA-seq analysis of non-fixed dissociated and fixed cell populations. Gene expression profiles were well correlated (*R* = 0.983), indicating that non-cross-linking fixation by CellCover could preserve the transcriptome profile of cells (Figure S1A).

**Figure 1:**
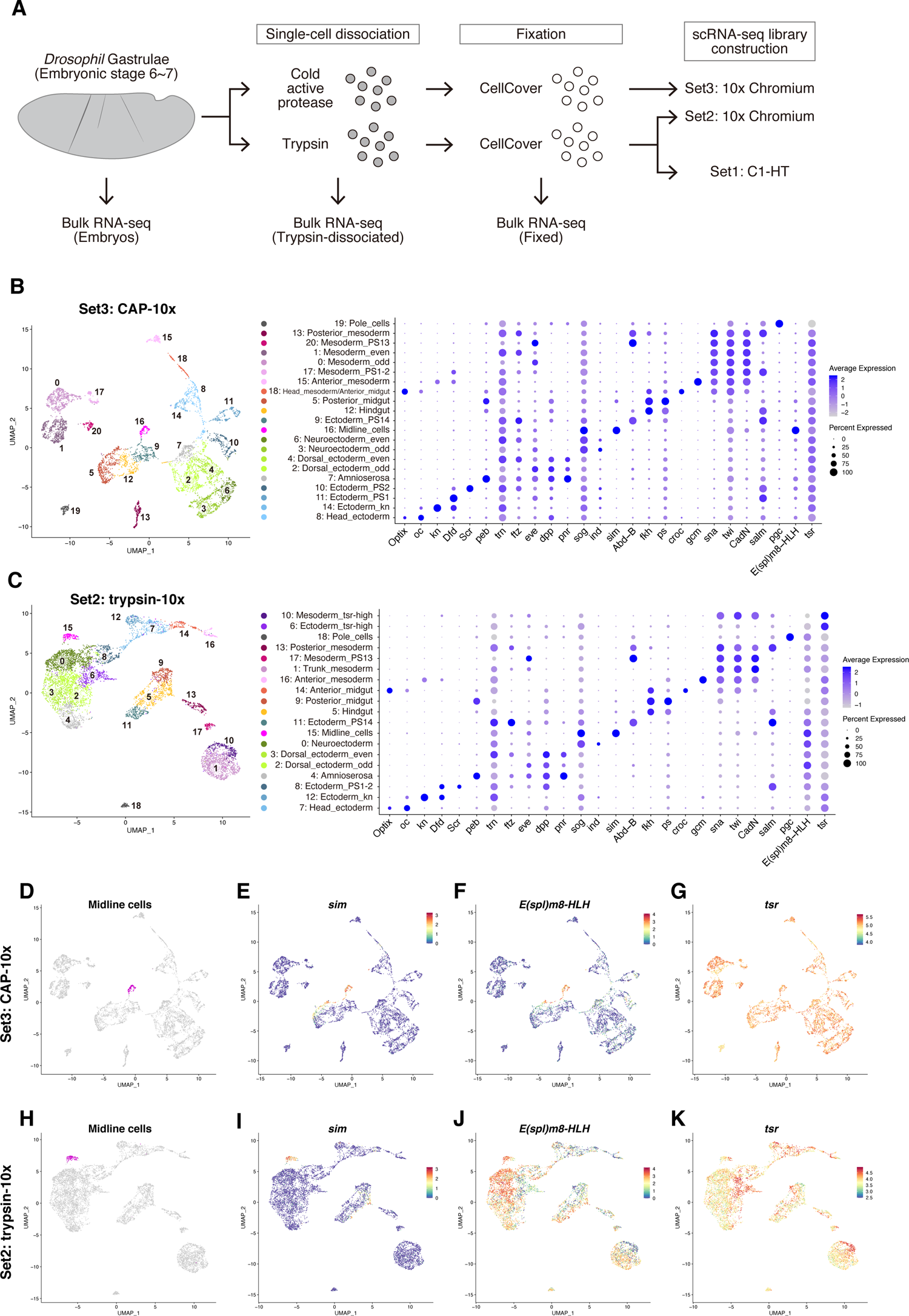
Comparison between trypsin and CAP dissociation for scRNA-seq. A. Schematic diagram of the data acquired in this study. B-C. Uniform manifold approximation and projection (UMAP) plot of the Set3 CAP-10x scRNA-seq data (B) and the Set2 trypsin-10x scRNA-seq data (C) with cluster information. Dot plot shows the expression patterns of typical marker genes for each cluster. D. Midline cells are colored magenta in the UMAP plot of the Set3 CAP-10x data. E-G. Expression patterns of *sim* (E)*, E(spl)m8-HLH* (F) and *tsr* (G) in Set3 CAP-10x data. H. Midline cells are colored magenta in the UMAP plot of the Set2 trypsin-10x data. I-K. Expression patterns of *sim* (I)*, E(spl)m8-HLH* (J) and *tsr* (K) in Set2 trypsin-10x data. The expression patterns in (E-G, I-K) represent the log-transformed values after SCTransform normalization.

We then performed scRNA-seq analysis using three different protocols: Set1 trypsin-dissociation and Fluidigm C1 mRNA Seq HT IFC (Set1 trypsin-C1HT), Set 2 trypsin-dissociation and 10x Genomics Chromium V3.1 (Set2 trypsin-10x), and Set 3 CAP-dissociation and 10x Genomics Chromium V3.1 (Set3 CAP-10x) (Figure 1A). In all dataset, gene expression was quantified by counting the different unique molecular identifiers (UMI), which were short random nucleotide sequences added to each transcript in the reverse transcription step, per gene and per cell (Islam et al., 2014). After filtering the data of high-quality cells (See Materials and Methods for details), 1,243, 7,314, or 6,180 cells remained and the expression of 4,500, 3,222, or 4,053 genes per cell in the median was detected for the Set1, Set2, or Set3 data, respectively. Set1 trypsin-C1HT data showed higher median UMI counts per cell (152,279) than the other 10x data (22,506 (Set2) and 37,610 (Set3)). One of the reasons for higher sensitivity is greater sequencing depth per cell, and this is consistent with a previous report that the C1 platform can produce rich information (Torre et al., 2018). Unsupervised clustering and the extraction of marker genes for each cluster using Seurat v3 (Stuart et al., 2019) revealed that each dataset contains all major cell types, such as mesoderm (*snail* (*sna*), *twist* (*twi*)), trunk dorsal ectoderm (*decapentaplegic* (*dpp*), *pannier* (*pnr*)), trunk neuroectoderm (*short gastrulation* (*sog*)), head ectoderm (*Optix*, *ocelliless* (*oc*)), terminal endoderm (*fork head* (*fkh*)), pole cells (*polar granule component* (*pgc*)), and dorsal amnioserosa cells (*pebbled* (*peb*)), indicating there was no cell type bias in all datasets (Figures 1B, C; Figure S1B). The Set1 trypsin-C1HT data was composed of four biological replicates, and there was no obvious batch effect among them, indicating the reproducibility of our protocol (Figure S1B). 10x data allowed us to identify more cell types than C1HT data, suggesting that sequencing more cells, rather than deeper sequencing of each cell, is more important for identifying minor cell types in an embryo that is composed of many cell types, as already mentioned (Heimberg et al., 2016; Zhang et al., 2020). The Set1 trypsin-C1HT data are useful for detecting low-expression genes and characterizing each cell more comprehensively.

### Trypsin-dissociation causes artificial upregulation of Notch-target genes

To reveal the extent to which different dissociation methods affect the single-cell transcriptome profile, we inspected the marker genes of each cluster for all scRNA-seq data in depth. Midline cells (mesoectoderm) are known to highly express Notch target genes, such as *single-minded* (*sim*) and some *Enhancer of split* (*E(spl)*) complex genes, such as *E(spl)m5, helix-loop-helix* (*E(spl)m5-HLH*), and *E(spl)m8-HLH* (Cowden and Levine, 2002; Morel and Schweisguth, 2000; Zinzen et al., 2006). In Set3 CAP-10x data, all these genes were identified as marker genes of the midline cell cluster (Figures 1D-F). On the other hand, we noticed that, in Set2 trypsin 10x data, although the midline cell cluster was identified by specific expression of *sim*, *E(spl)m8-HLH* showed strong and broad expression not only in the midline cell cluster, but also in other clusters (Figures 1H-J). In the cell dissociation process from embryo homogenization to cell fixation, cells were placed on ice except for 10 min trypsin treatment, suggesting that trypsin treatment caused the upregulation of *E(spl)* complex genes through artificial activation of Notch signaling irrespective of cell type. To assess whether artificial induction of *E(spl)* complex genes is due to trypsin treatment, we analyzed bulk RNA-seq data to identify differentially expressed genes (DEGs) between intact embryos and trypsin-dissociated cells. Although there was a high correlation between them, trypsin-dissociated cells showed higher expression of some *E(spl)* complex genes, indicating that they were artificially upregulated by trypsin treatment before fixation (Figures S2A, B).

In addition, Set2 trypsin-10x data had clusters that showed high *twinstar* (*tsr*) expression (Figure 1K). For example, the trunk mesoderm was divided into two clusters (clusters 1 and 10 in Figure 1C), and cluster 10 showed higher expression of *tsr* compared to cluster 1 (Figure 1K). Furthermore, cluster 6, which seems to belong to the trunk ectoderm, also showed high *tsr* expression. Clusters 6 and 10 showed relatively low expression of *E(spl)* complex genes (Figure 1J). To characterize these cells with high *tsr* expression (*tsr*-high cells), we identified highly expressed genes in cluster 10 compared to cluster 1 and performed gene-set enrichment analysis. By Gene Ontology (GO) enrichment analysis, we found that the term “oxidative phosphorylation” was enriched in highly expressed genes of the *tsr*-high cell cluster (Figure S2C), suggesting that these cells exhibited some metabolic stress responses.

These results suggest that there are two types of cells in Set2 trypsin-10x data: one increased some of the Notch target genes (*E(spl)* complex genes), and the other showed some kind of stress response upon trypsin treatment. Set1 trypsin-C1HT data also showed strong and broad expression of *E(spl)* complex genes (Figures S1D, E), and clusters showing high *tsr* expression (Figure S1F). On the other hand, in Set3 CAP-10x data, there was no such cluster (Figures 1F, G), indicating that these cellular responses are specific to trypsin treatment, but not CAP treatment. Notably, even in Set2 trypsin-10x data, similar clusters were distinguished as in Set3 data (Figure 1C), indicating that these trypsin-dependent artifacts did not extensively distort the transcriptome profile. Because it is likely that the CAP dissociation could well preserve the original expression patterns of Notch target genes, and there was no cluster showing any stress responses, we mainly focused on the Set3 CAP-10x dataset using CAP for further analysis.

### Identification of 65 transcriptomically distinct clusters

To further investigate the detailed single-cell transcriptome diversity in the gastrulae, we focused on the Set3 data and manually annotated each cell utilizing known gene expression patterns from databases (BDGP *insitu* database (https://insitu.fruitfly.org), Fly-FISH (http://fly-fish.ccbr.utoronto.ca)), and information from the literature. In the trunk region of the gastrula, the amnioserosa, dorsal ectoderm, ventral neuroectoderm, mesoectoderm (midline cells), and mesoderm emerge along the dorsal-ventral axis (Reeves and Stathopoulos, 2009). On the other hand, along the anterior-posterior axis, cells were divided into 14 parasegments (PS) (Akam, 1987; Ingham, 1988; Jaeger, 2011; Jaeger et al., 2012; McGinnis and Krumlauf, 1992; Peter A. Lawrence, 1992), and even parasegments express *tartan* (*trn*) and *ftz* specifically (Clark and Akam, 2016; Graham et al., 2019). Unsupervised graph-based clustering with 3,000 highly variable genes (HVGs) revealed that cells in the trunk ectoderm were divided into mixed characteristics of both AP- and DV-axis patterns (Figure 1B. clusters 2, 3, 4 and 6). For example, cluster 2 was the dorsal part of odd parasegments, and cluster 6 was the ventral neuroectoderm of even parasegments. To annotate the spatial origin more precisely, we inferred the origin of the cells for each of the AP and DV axes separately. We picked up the trunk ectoderm cells (Figure 2A, corresponding to PS2-13), and assigned these cells to seven DV identities (Amnioserosa (*zerknullt* (*zen*) +, *peb*+), medial dorsal ectoderm (*eiger* (*egr*) +, *Dorsocross2* (*Doc2*) +, *zen*–), intermediate dorsal ectoderm (*dpp*+, *extra macrochaetae* (*emc*) +, *Doc2*–), lateral dorsal ectoderm (*dpp*+, *Ataxin 1* (*Atx-1*) +), lateral neuroectoderm (*Drop* (*Dr*) +, *SoxNeuro* (*SoxN*) +), intermediate neuroectoderm (*intermediate neuroblasts defective* (*ind*) +, *SoxN*+, *sog*+), medial neuroectoderm (*ventral nervous system defective* (*vnd*) +, *SoxN*+, *sog*+)) by k-means clustering with 35 selected DV genes (Figure 2D). Along the AP axis, we annotated each cell based on graph-based clustering by Seurat and Hox gene expression as landmarks, and assigned these cells to four AP identities (Parasegment 2 (PS2), Trunk ectoderm odd (PS3, 5, 7, 8, 9, 11), trunk ectoderm (PS4, 6, 8, 10, 12), and PS13) (Figures 2B, C). By combining these AP and DV identities, the trunk ectoderm cells were divided into 25 subclusters (Figure 2E). We also performed a manual annotation for each cluster (Figure S3), and eventually divided the cells into 65 subclusters for Set3 data (Figure 3; Figure S4). The mesoderm was divided into 14 types along the AP axis (Figures S3A-E). Posterior clusters (clusters 5, 9, and 12) were composed of three posterior midgut types, two hindgut types, and four types of parasegment 14 ectoderm (Figure S3H). Head regions located anterior to parasegment 1 could be divided into two anterior endoderm and seven head ectoderms containing future foregut primordium (Figures S3F, G). The results of the subclustering are summarized in Supplementary Table 1. During the subclustering process, 62 potential doublet cells were identified and discarded from the dataset. The remaining dataset consisted of 6,118 cells. We noticed that two of the 65 clusters were difficult to annotate with an equivocal identity. From the subclustering of cluster 18, three subclusters were identified (Figure S3B). One of them was “Anterior endoderm wg+” that specifically expresses anterior endoderm markers, such as *fkh*, *huckebein* (*hkb*), and *serpent* (*srp*), as well as *wg*. Another subcluster named “Anterior mesoderm” showed the expression of mesoderm markers, such as *sna*, *twi*, and *heartless* (*htl*). However, the third subcluster expressing *wnt inhibitor of Dorsal* (*wntD*) was positive for both endoderm and mesoderm markers. Since *wntD* is known to repress mesoderm differentiation (Ganguly et al., 2005; Rahimi et al., 2016), we annotated this subcluster as “Anterior endoderm wntD” at this moment. Another example of an intermediate state is “PS14 ectoderm wg” which seems to be between PS14 ectoderm and hindgut (Figure S3H). It expressed both PS14 ectoderm markers (*Abdominal B* (*Abd-B*)) and hindgut markers (*disconnected* (*disco*), *wg*, and *no ocelli* (*noc*)). Since this subcluster did not show the expression of the key hindgut gene *fkh*, we annotated this subcluster as one of the PS14 ectoderm regions. We concluded that the scRNA-seq data contains enough information, in terms of both the number of cells and sequencing depth, which allows us to distinguish the spatial origin at the single-cell level, as well as rare intermediate states that have not been recognized so far.

**Figure 2:**
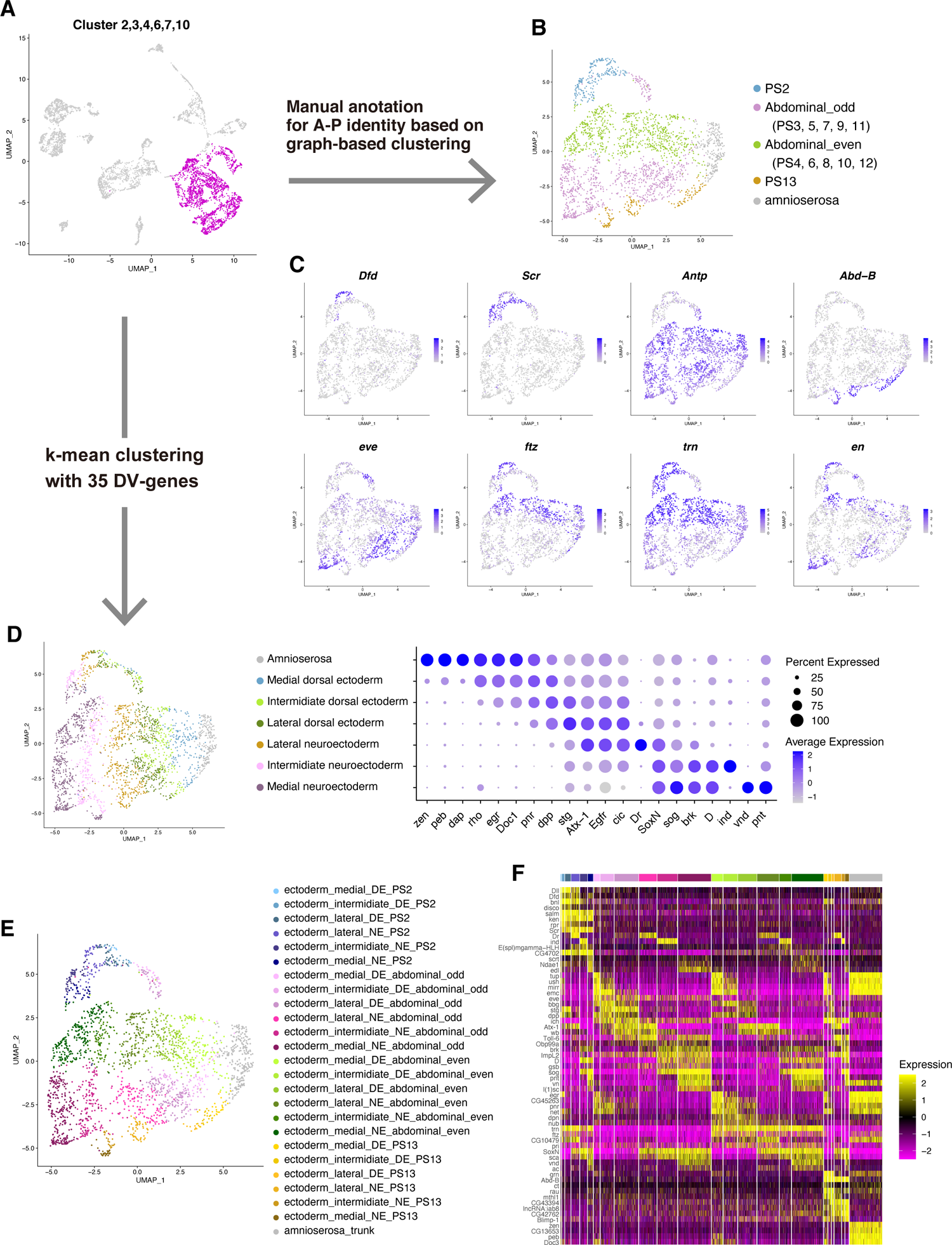
Subclustering of trunk ectodermal cells. A. Trunk ectodermal cells are colored magenta in the UMAP plot of the Set3 CAP-10x data. B. Annotation of the anterior-posterior (AP) identities of the trunk ectodermal cells. C. Expression patterns of anterior-posterior specific genes. D. Annotation of dorsal-ventral (DV) identities of the trunk ectodermal cells. Dot plot shows the expression patterns of typical marker genes for each cluster. E. Annotation of the trunk ectodermal cells with a combination of AP and DV identities. F. Heatmap showing the typical marker genes for all subclusters of the trunk ectodermal cells.

**Figure 3:**
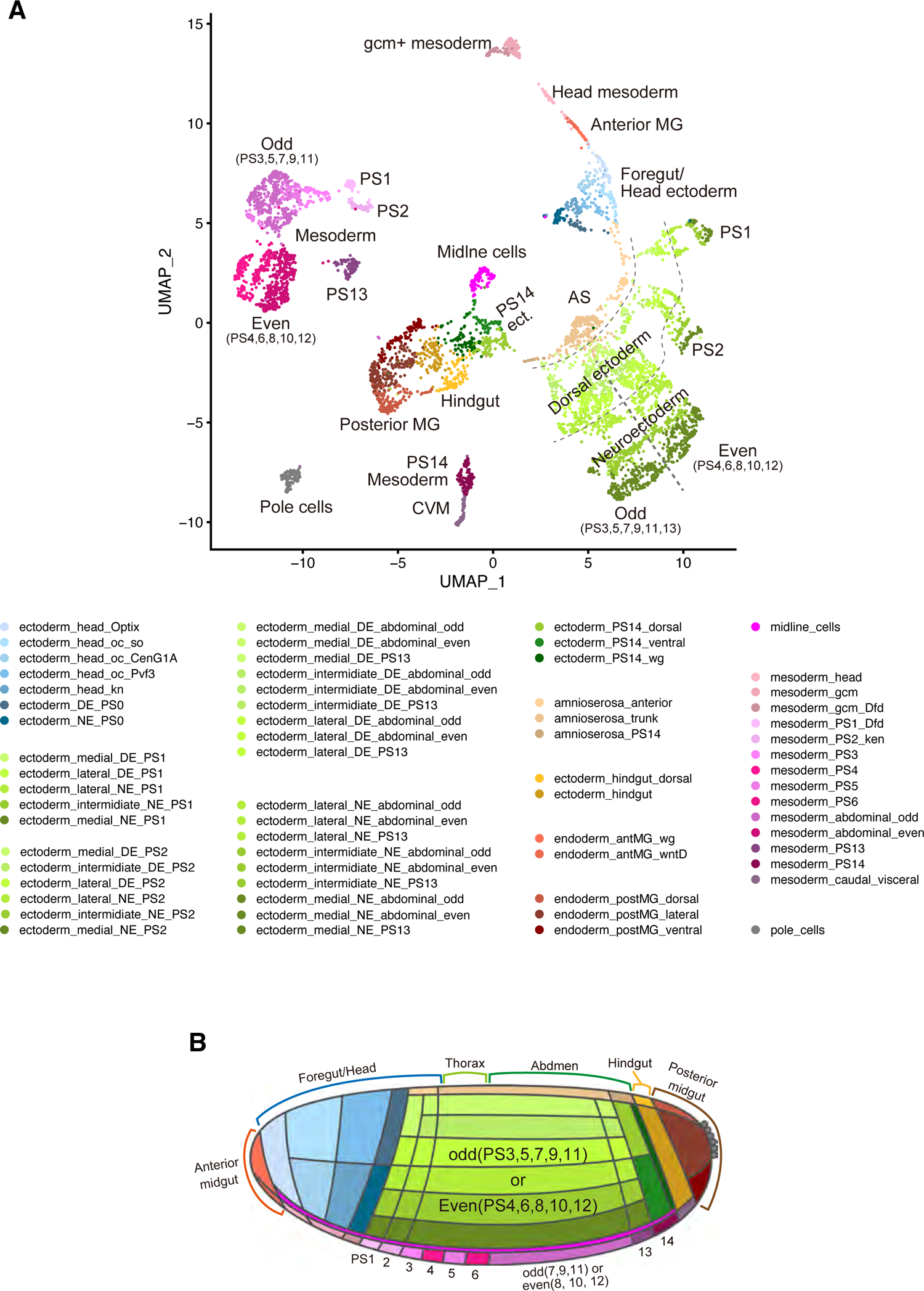
65 subclusters of *Drosophila* gastrulae identified from scRNA-seq data. A. UMAP plot of the Set3 CAP-10x scRNA-seq data with information on the 65 subclusters. B. Schematic diagram showing the inferred spatial location of each subcluster in gastrulae. AS: Amnioserosa, CVM: Caudal visceral mesoderm, DE: Dorsal ectoderm, MG: Midgut, NE: Neuroectoderm

### Expression profile of plasma membrane genes, rather than transcriptional regulator genes, better represents the major cell types

Next, we analyzed the features of the transcriptome profile that contributed to the classification of each cell. GO term analysis revealed that genes encoding transcriptional factors (TF genes), as well as plasma membrane-related genes (PM genes) were highly enriched in 1,500 HVGs (Figures 4A, B). This suggests that a combination of transcription factors and downstream effector plasma membrane-related genes mainly contributes to the generation of transcriptome diversity in *Drosophila* gastrula. Hierarchical clustering of 64 subclusters (pole cells were removed) with all 1,500 HVGs classified the subclusters into three germ layers (ectoderm, endoderm, and mesoderm), indicating that the differences in the cellular transcriptome at the gastrula stage reflect the differences among future cell lineages (Figures 4C, F). However, it is unclear whether each gene set alone holds enough information to characterize the cell type. To address this issue, based on the Gene List Annotation for *Drosophila* database (GLAD; https://www.flyrnai.org/tools/glad/web/) (Hu et al., 2015), the 1,500 HVGs were classified into four categories (TF genes, PM-related genes, non-TF and non-PM genes in GLAD, and genes not included in GLAD; see Materials and Methods for details)(Figure S5A), and clustering analyses were performed using each gene set alone. The set of 258 TF genes in 1,500 HVGs well segregated each cell along with the original positions on the UMAP plot (Figure 4D). By hierarchical clustering using only TF genes, subclusters tended to be classified by their spatial location compared to the case using all 1,500 HVGs (Figure 4G). For example, the subclusters of “ectoderm_PS14”, “mesoderm_PS14” and “mesoderm_caudal_visceral” form a single group beyond the types of mesoderm and ectoderm. In addition, the subclusters located in anterior and posterior terminals were also grouped across the three germ layers. On the other hand, the set of PM genes well reproduced the clustering pattern with all 1,500 HVGs, and hierarchical clustering categorized 64 subclusters with their germ layer identities (Figures 4E, H). Furthermore, the set of non-TF and/or non-PM genes also classified the subclusters into three germ layers (Figures S5B-E). These clustering analyses revealed that, without any prior functional knowledge about each gene, only the mRNA expression profiles of TF genes were insufficient to distinguish future cell lineages. On the other hand, those of other effector genes better represented the differentiation status of the three germ layers.

**Figure 4:**
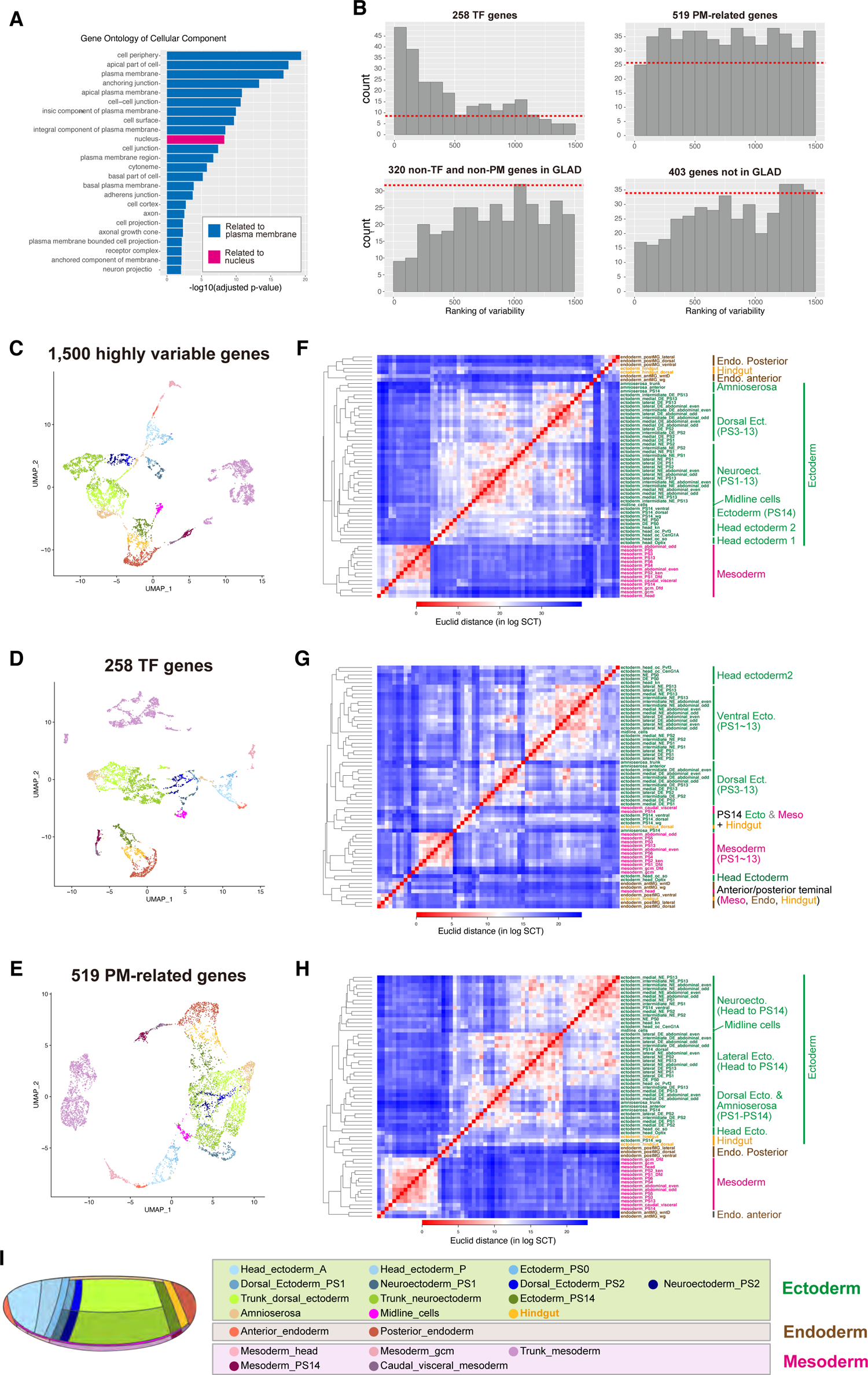
Clustering analysis with GLAD categories. A. Gene Ontology enrichment analysis of 1,500 HVGs using g:Profiler. The terms of cellular components are presented. B. The number of genes belonging to each category in each bin of 1,500 HVGs divided into 15 bins from the top. The red dotted line is the expected number by random sampling. C-E. UMAP plot with 1,500 HVGs (C), 258 TFs in 1,500 HVGs (D), and 519 PM-related genes in 1,500 HVGs (E). Cells were colored by super cluster information based on prior annotations (I) F-H. Hierarchical clustering analyses of 64 subclusters based on the Euclidean distances in log-transformed gene-expression space with top 1,500 HVGs (F), 258 TFs in 1,500 HVGs (G), and 519 PM-related genes in 1,500 HVGs (H). Subclusters were colored based on the future germ layers. I. Super cluster information used to color the cells in the UMAP plot. See Supplementary Table S1 for details.

### Transcriptome-level differences between the single-cell stripes along the A-P axis

During gastrulation of *Drosophila* gastrulae, each parasegment of the lateral ectoderm is composed of four stripes along the A-P axis, and each stripe is a single-cell wide column (Figure 5A). Each stripe has different identities with combinatorial sets of pair-rule genes, directing polarized myosin localization and cell intercalation movement (Bertet et al., 2004; Zallen and Blankenship, 2008; Zallen and Wieschaus, 2004). Because pair-rule genes encode TFs, they should control downstream effector genes to drive germband extension. Some effector genes, such as *18 wheeler* (*18w,* also known as *Toll-2*), *Toll-6* and *Tollo* (also known as *Toll-8*), and *trn* were identified (Paré et al., 2014, 2019), but the sufficient conditions for differential gene expression to promote germband extension are still unclear. For example, there is a possibility that several different mechanisms act in parallel to drive germband extension, and minor mechanisms may have been overlooked due to the large contribution of major mechanisms. In addition, it has been proposed that, from detailed live imaging analyses, the difference between the third and fourth stripes in each parasegment might be difficult to distinguish, suggesting that the strength of cell-cell interaction between them is weaker and the difference in gene expression profiles may also be smaller (Tetley et al., 2016). Tetley et al. also proposed that ‘super-boundaries’ that interface between cells of non-adjacent stripes (‘skipped boundary’) are more contractile, implying that these boundaries have larger differences in receptor expression patterns and stronger cell-cell interaction than boundaries of adjacent identities (Tetley et al., 2016).

**Figure 5:**
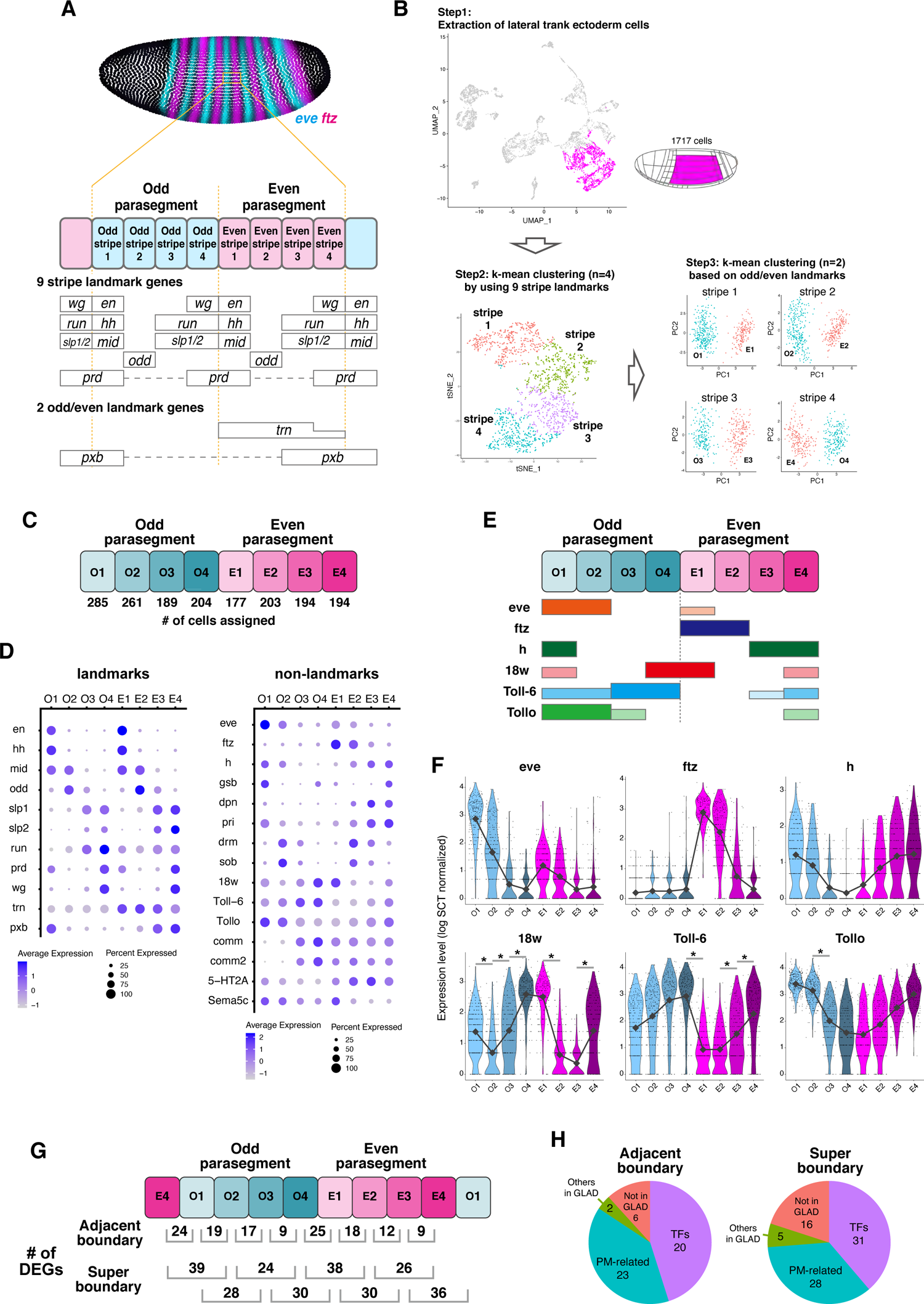
Assignment of the stripe identities to ectodermal cells. A. (top) Examples of the stripe expression patterns of pair-rule genes (*eve*, *ftz*) from the BDTNP ISH database. *eve* is expressed in odd parasegments and *ftz* is expressed in even parasegments. (bottom) Expression patterns of nine stripe landmark genes and two odd/even landmark genes. B. (Step 1) UMAP plot of the Set3 CAP-10x data. Trunk ectodermal cells used for stripe assignment are colored magenta. (Step 2) t-SNE plot with 1,711 trunk ectodermal cells and 9 stripe landmark genes. Cells were colored by k-means clustering (n=4) using 9 stripe landmark genes. (Step 3) Principal component analysis (PCA) plots of each stripe in Step 2. Cells were colored by k-means clustering (n=2) using *trn* for stripe 1 and 2 (or *pxb* for stripe 3 and 4) and genes positively and negatively correlated with it. C. The number of cells assigned to each stripe identity. D. Dot plots showing the expression patterns of landmark genes used for stripe assignment and non-landmark genes in each stripe. E. Reported stripe patterns of *eve*, *ftz*, *h*, *18w*, *Toll-6*, and *Tollo* (Clark and Akam, 2016, Paré et al., 2019). F. Violin plots showing the expression patterns of *eve*, *ftz*, *h*, *18w*, *Toll-6*, and *Tollo* in each reconstructed pair-rule stripe. The gray line indicates the median values for each stripe. Expression levels represent the log-transformed values after SCTransform normalization. G. The number of differentially expressed genes (DEGs) between adjacent or super boundaries. DEGs were identified using the FindMarkers function with MAST at a FDR threshold of 0.01 and FC threshold of 0.25. H. GLAD category breakdown for unique DEGs of adjacent or super boundaries.

To clarify the whole picture, we need to quantitatively describe the transcriptome of each stripe and understand how the gene expression profiles differ among them. To do this, by using pair-rule genes and segment polarity genes as stripe landmarks, we categorized the trunk ectodermal cells into eight single-cell stripes that span an odd- and even parasegmental unit (Figures 5A-C) (see Materials and Methods for details). To validate this striped pattern inference, we visualized the repeated pattern of genes that are already known to show striped expression along the A-P axis (Figure 5D). These visualizations revealed that, in addition to the landmark genes that were used for the inference, other stripe genes, such as *18w, Toll-6,* and *Tollo*, showed gradual changes, and these patterns were well correlated with the expression patterns revealed by *in situ* hybridization (Figures 5E, F) (Paré et al., 2019). These results indicated that the reconstruction of the stripe pattern was highly accurate.

This transcriptome information of eight single-cell stripes provides opportunities to quantitatively compare the differences in gene expression profiles between them. First, we conducted the DEG analyses between all pairs of adjacent identities. Based on an expression difference of ≥ 1.3 fold and a false discovery rate (FDR) of ≤ 0.01, nine to 25 genes were identified as DEGs between adjacent pairs (Figure 5G). Next, we investigated the DEGs. Similar to the DEG composition of whole scRNA-seq data, most of the DEGs between adjacent stripes were TF- or PM-genes, and there was little contribution from other cytoplasmic genes (Figure 5H). As proposed by the “Toll receptor code,” all boundaries showed at least one Toll receptor gene (*18w*, *Toll-6*, and *Tollo*) as DEGs (Figure 5F). *trn* was also differentially expressed at the parasegment boundaries. In addition to these known PM genes, our scRNA-seq data revealed that transmembrane genes, such as *commissureless* (*comm*), *comm2*, and *Semaphorin 5c* (*Sema5c*), were quantitatively differentially expressed in a stripe manner (Figure S6A), suggesting that these genes also play a role in cell-cell recognition for cell intercalation. In terms of the number of DEGs, the difference between parasegments (Odd4 vs. Even1 and Even4 vs. Odd1) was larger than that between cell stripes within parasegments, and the difference between the third and fourth stripes (Odd3 vs. Odd4 and Even3 vs. Even4) was the lowest (Figure 5G). In addition, comparing the differences between super-boundaries, more DEGs were identified than the differences between adjacent pairs (Figures 5G, H; Figure S6B). These results are consistent with the proposed models of super-boundaries and smaller differences between the third and fourth stripes (Tetley et al., 2016). This quantitative dataset will be important for determining the sufficient mechanisms for germband extension.

### scRNA-seq analysis of the *bicoid* mutant

During development, the input signals direct cell fate through gene regulatory networks. Perturbations of upstream signals often compromise the process and result in the transformation of one cell type to another. However, these transformations have been assessed by the change in the expression of limited marker genes, most of which encode TFs, and it is not clear whether the cells transformed at the level of the whole transcriptome. To address this issue, we performed scRNA-seq analysis of the *bicoid*-depleted embryos. The A-P axis of *Drosophila* is determined by the morphogen gradients of the anterior Bicoid (Bcd) and the posterior Nanos (Nos). It is widely accepted that the loss of Bcd function results in the conversion of the anterior identity into the posterior one (Petkova et al., 2019; Staller et al., 2015). This transformation has been demonstrated by a limited number of gene expression and morphogenetic phenotypes, and it is still unclear whether this transformation occurs at the transcriptome level. The anterior part of *bcd* mutants eventually shows posterior profiles, but the developmental history of cells in the anterior region to reach the state is different from that in the original posterior region. For example, the onset of the anterior *hunchback* (*hb*) expression in *bcd* mutants is delayed compared to that in the posterior (Staller et al., 2015). In addition, pair-rule genes *ftz* and *eve* show mutually exclusive expression patterns of each other in wildtype, but they are initially overlapped in the early stage of *bcd* mutants, and then segregated later (Staller et al., 2015). Since these historical differences may affect the final state (Briscoe and Small, 2015), it is still possible that there are transformed cells with a mixed state of both anterior and posterior identities at the transcriptome level.

To clarify these points, we conducted scRNA-seq of *bcd*-RNAi (*bcd*-depleted) embryos and compared the cellular composition and transcriptome with that of control embryos. In *bcd*-RNAi embryos, cell types that belong to the anterior region of wild-type embryos, such as the anterior midgut, head/PS1-2 ectoderm, and anterior mesoderm were not identified (Figures 6A, B). When the *bcd*-RNAi data were integrated with the control data, there were no novel cell types or no mis-specified cells in *bcd*-RNAi embryos (Figure 6C). The anterior clusters consisted of control cells only (Figure 6C, Head_ectoderm, Ectoderm_PS1-2, anterior midgut/mesoderm). The ratio of cells assigned to posterior clusters in *bcd*-depleted embryos was almost double that in wild-type embryos (Figure 6D). These results indicate that the anterior region of *bcd*-RNAi embryos is completely canalized into the posterior identity, and there are no mixed-state cells at the transcriptome level. We consider that this complete transformation could be achieved without posterior determinants because the *bcd* mutation does not affect the mRNA localization and translation of the posterior determinant *nanos* (Wang et al., 1994). This complete transformation of the transcriptome is consistent with the observation that the anterior region of *bcd* mutants exhibits the gastrulation movement of the posterior pole, as well as its differentiation to posterior identity. Similar results have been reported in scRNA-seq of zebrafish mutants with defective Nodal signaling (Farrell et al., 2018; Wagner et al., 2018). Taken together, these results strongly suggest that there are strict constraints in gene regulatory networks that allow cells to canalize into the defined transcriptomic state existing in wild types upon perturbations.

**Figure 6:**
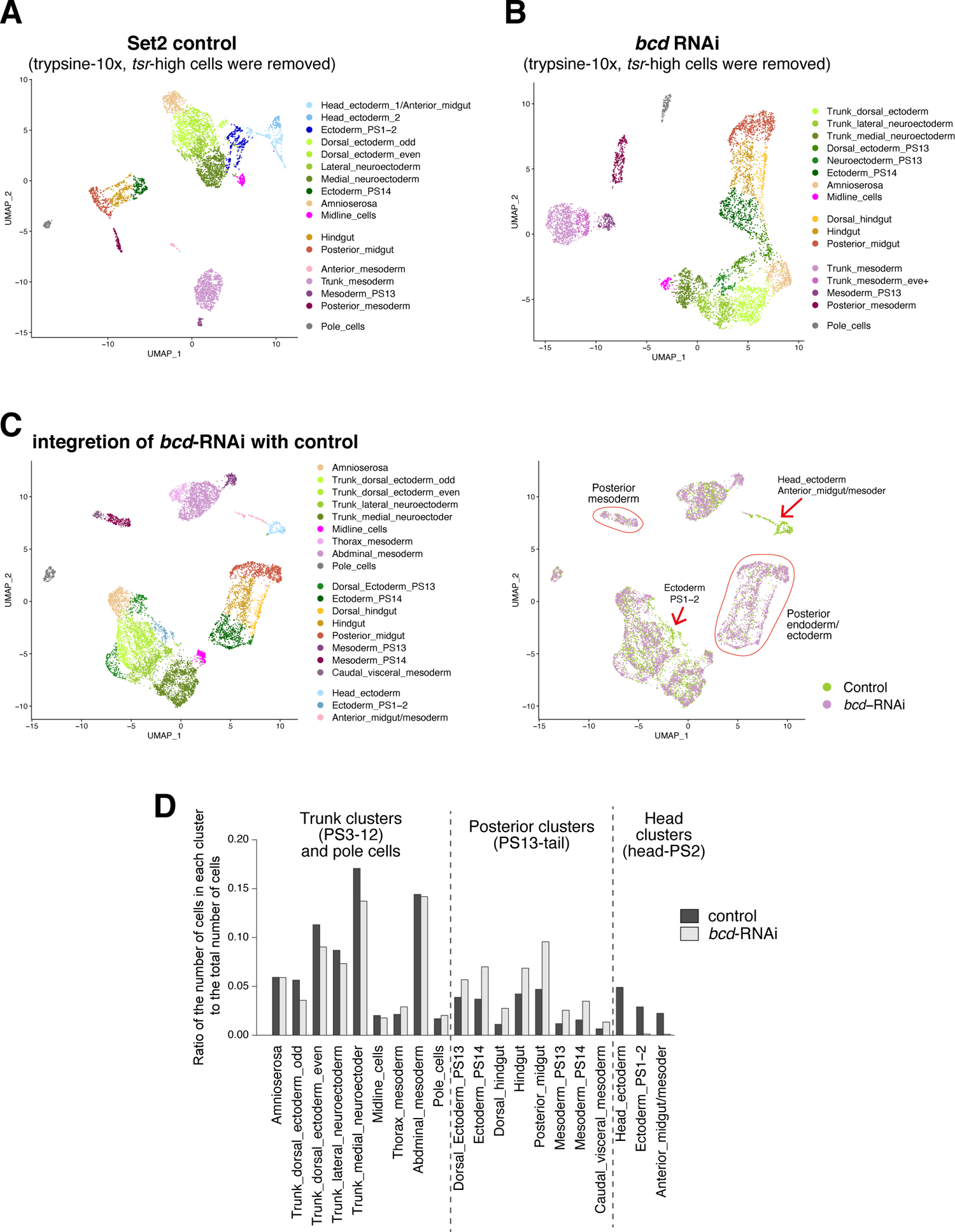
Analysis of fate transformation in *bcd* knockdown mutants with scRNA-seq. A. UMAP plot of Set2 trypsin-10x data after removal of *tsr*-high cells. B. UMAP plot of *bcd*-KD scRNA-seq data after removal of *tsr*-high cells. C. (left) UMAP plot of integrated data between Set2 (A) and *bcd*-KD (B). Clustering and annotation were performed again after integration. (right) Cells are colored according to the data they were derived from. Arrows indicate the clusters that are only composed of cells from the Set2 control. D. Ratio of the total number of cells to the number of cells in each cluster for each dataset.

### Spatial reconstruction of all gene expression patterns at the single cell resolution

We and others have made efforts to reconstruct the spatial expression patterns of all genes at single cell resolution by developing novel computational tools. However, all reconstructions were based on the scRNA-seq data composed of 1,297 cell data established by N. Karaiskos et al. (hereafter, NK-data)(Karaiskos et al., 2017), and the number of cells was much smaller than that in a *Drosophila* gastrula. Here, we obtained new scRNA-seq data containing 6,118 cell data (hereafter, Set3), which is approximately equal to the number of cells in a *Drosophila* gastrula. We hypothesized that our Set3 data would substantially improve the reconstruction quality.

We reconstructed the spatial expression pattern using our Set3 data and Perler (Okochi et al., 2021), and then compared the results with those obtained using the NK-data. The reconstruction method uses quantitative *in situ* hybridization (ISH) data from BDTNP as a reference for spatial patterns. However, both scRNA-seq data were derived from stage 6-7 embryos, while ISH data were established for stage 5 embryos. We noticed that some of the genes in the ISH data dynamically changed the expression pattern from stage 5 to stage 6-7. Therefore, we removed the 26 genes from the reference, whose expression patterns significantly changed between the two time points and could worsen the reconstruction results; hence, 67 genes were used as references (Supplementary Table S6). To compare Set3- and NK-data-based reconstruction using Perler, we first performed leave-one-gene-out cross validation (LOOCV) for both datasets. Set3-based reconstruction showed a higher prediction score (median correlation coefficient = 0.66) than that of NK-data-based reconstruction (median correlation coefficient = 0.61) (Figure 7A), indicating that the Set3-based reconstruction had a higher generalization performance than the NK-data-based reconstruction. Second, the gene-gene correlations in scRNA-seq data were better conserved in Set3-based reconstruction than in NK-data-based reconstruction (Figure 7B). Finally, Set3-based reconstruction maintained the scale of expression values, but NK-data-based reconstruction did not (Figures 7C-E). For example, NK-data-based reconstruction had high background values and had to be rescaled by the minimum to maximum value for each gene. On the other hand, Set3-based reconstruction showed quite low background signals, and there was no need to perform any re-scaling. The expression values in Set3-based reconstruction were comparable to those in the original scRNA-seq data. In the spatial reconstruction based on NK-data, maybe due to the insufficient number of sequenced cells, a given reconstructed cell is likely to have more contribution from less related scRNA-seq data.

**Figure 7:**
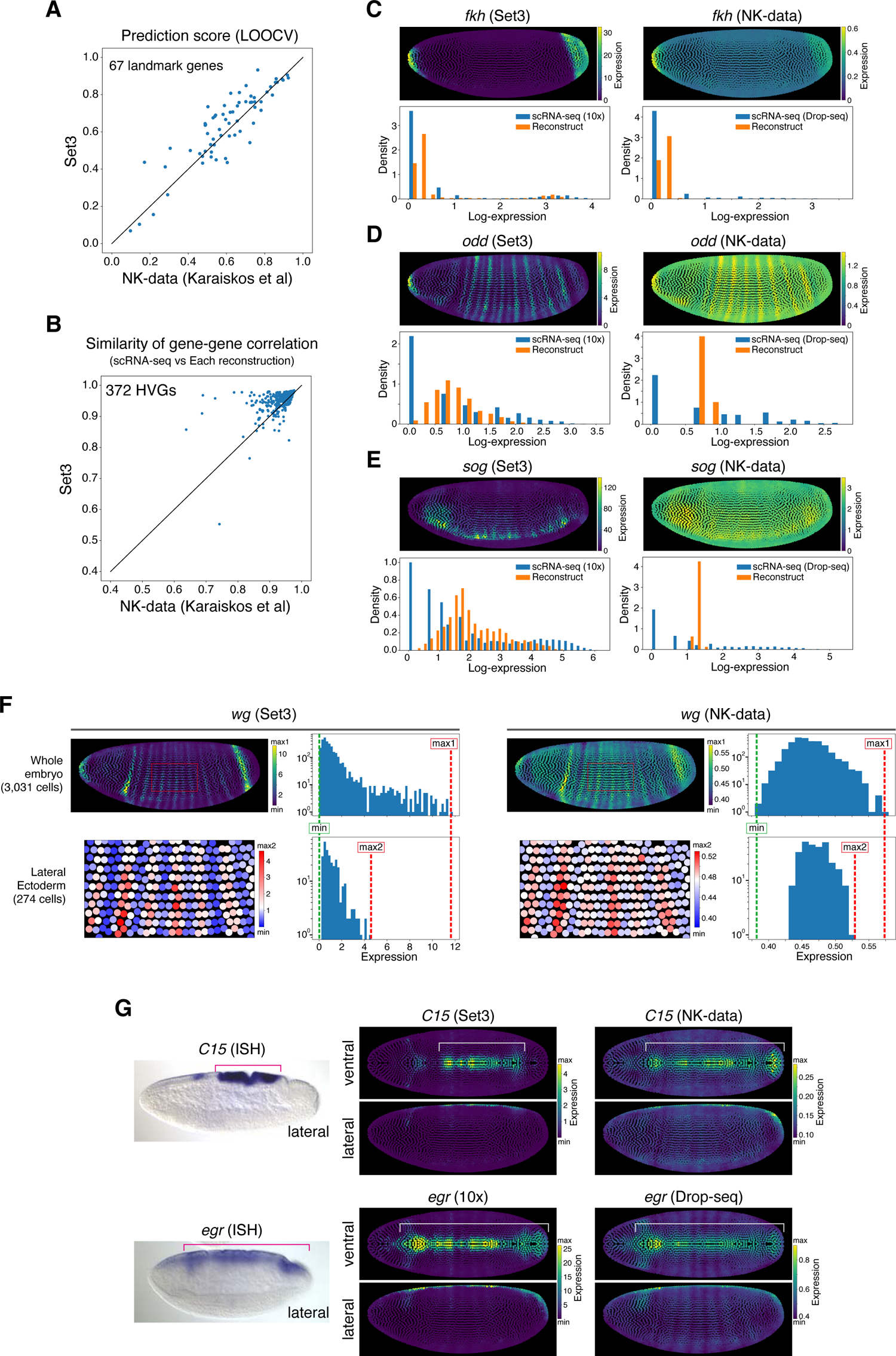
Spatial reconstruction by Perler. A. Comparison between the leave-one-gene-out cross validation (LOOCV) results of Perler reconstructions based on NK-data and Set3 data. Each dot indicates each landmark gene. X- and Y-axis show the correlations between reference ISH expression patterns and the reconstructed expression pattern of each gene based on NK-data and Set3 data, respectively. B. Comparison of gene-gene correlation structure conservation between Set3-based and NK-data-based reconstruction by Perler. Each dot indicates each gene which was commonly included in Top 500 HVGs of both datasets (372 genes). The X-axis shows gene-gene correlation structure conservation in NK-data-based reconstruction and the Y-axis shows gene-gene correlation structure conservation of Set3-based reconstruction. The definition of gene-gene correlation structure conservation was described in Materials and Methods. C-E. Examples of reconstructed expression by Perler on Set3 (left) and NK-data (right). In each plot, upper panels show reconstructed expression patterns. Colormaps are linear and zero-max scaled. Bottom panels show density histograms of the gene expression in the original scRNA-seq and the reconstruction. Expression patterns are log-scaled and each bin size is 0.2 in density histograms. F. The reconstructed expression patterns of *wg* by Perler based on Set3 (left) and NK-data (right). In each plot, upper-left panel shows reconstructed expression patterns in whole-embryo. Bottom-left panel shows the enlarged views of the region enclosed with a red rectangle in the upper-left panel. Upper-right panel and bottom-right panel show the histograms of the expression in whole-embryo and the region shown in bottom-left panel, respectively. Expression values in plots are linear-scaled and on y-axes are log-scaled in the histogram. The maximum expression patterns in the whole embryo plots and enlarged plots are indicated by “max1” and “max2” (red dashed lines), respectively. The minimum expression in whole embryo is indicated by “min” (green dashed line) G. (left) *in situ* hybridizations of *C15* and *egr* from the Berkeley Drosophila Genome Project (BDGP; https://insitu.fruitfly.org/) (Hammonds et al., 2013; Tomancak et al., 2002, 2007). Lateral view. (middle) The lateral or dorsal views of Set3-based reconstructed patterns of *C15* and *egr* by Perler. (left) The lateral or dorsal views of NK-data-based reconstructed patterns of *C15* and *egr* by Perler. Red and white lines show the region of expression along the dorsal midline.

We also found that the reconstructed pattern of some genes was qualitatively improved using our Set3 data. For example, the pattern of segment polarity genes (*wg* and *en*) became much clearer in Set3 reconstruction. NK-data-based reconstruction showed weak *wg* expression in the cells between the stripes, while this background was suppressed to almost zero in the Set3-based reconstruction (Figure 7F). In addition, ISH for *C15* showed expression only in the dorsal amnioserosa, and ISH for *egr* showed a broader expression along the dorsal midline. The reconstructed pattern of *C15* with NK-data and Perler showed a broad expression like *egr*, while Set3 and Perler reconstructed a pattern similar to that of ISH (Figure 7G). These results indicate that the reconstruction based on Set3 data is more accurate and provides better interpretability for applications in future biological studies.

Recently, in addition to Perler, other computational methods for spatial reconstruction have been proposed. One of them is NovoSpaRc, which adopts a different strategy from Perler, and is based on the hypothesis that physically neighboring cells share similar transcriptional profiles and the framework of optimal transport (Moriel et al., 2021; Nitzan et al., 2019). To compare their performances when our Set3 data were used as input and to establish a more accurate reconstruction, we attempted spatial reconstruction using NovoSpaRc and our Set3 data. Overall, both Perler and NovoSpaRc showed comparable performances. First, the spatial reconstruction by Perler and NovoSpaRc showed a high correlation with each other (Figure S7A). Second, the prediction performance of spatial reconstruction by LOOCV were also comparable to each other (Figure S7B). Finally, we also examined the degree to which the spatial reconstruction by Perler and NovoSpaRc conserved the gene-gene correlation in the original scRNA-seq, and found that Perler maintained slightly higher gene-gene correlations than NovoSpaRc (Figure S7C). On the other hand, from a qualitative point of view, NovoSpaRc showed more spatially uniform patterns than Perler. For example, both methods well reconstructed the ventral expression of mesodermal gene *twi*, the expression within ventral mesodermal in Perler looked more spatially variable than that in NovoSpaRc (Figure S7D). This could be due to the fact that NovoSpaRc takes physical distances between cells into account. The reconstructed patterns obtained using both methods are listed in Supplementary Figures S8 and S9.

Although both reconstructions seem to be highly accurate, there is still a limitation. In both methods, the spatial gene expression along the anterior-posterior axis appears to be well reconstructed, whereas that along the dorsal-ventral axis seems to be insufficient. For example, in the UMAP plot of the original scRNA-seq data, the expression levels of *vnd* and *ind* were mutually exclusive in the *brk*+ medial neuroectoderm, intermediate neuroectoderm, and midline cells, but these levels were intermingled in both reconstructed data (Figures S7E-H). This is probably because the reference BDTNP ISH data is not sufficient and accurate because of the limited number of genes analyzed and the incomplete computational integration of the imaging data from multiple embryos. The construction of more precise reference data in the future will enable us to perform a more accurate reconstruction of the spatial transcriptome. Taken together, at this moment, it would be better to use a combination of both reconstruction results with the original scRNA-seq data in future biological applications.

## Discussion

We conducted scRNA-seq analysis to establish the single-cell transcriptome atlas of *Drosophila* gastrulae with higher accuracy and spatial resolution. This data consists of 6,118 cells that cover the entire gastrula, and allowed us to identify 65 subclusters. We also recapitulated the stripe expression patterns along the A-P axis with single-cell wide column resolution. We found that, at the transcriptome level, rather than the primary TF layer in the regulatory network, the subsequent layer of PM-related genes or other cytoplasmic genes showed mRNA expression profiles that better represented the features of the three germ layers. A new spatially reconstructed dataset is also established.

### Artificial effect of trypsin treatment during cell dissociation

Single-cell dissociation is one of the critical steps for scRNA-seq analysis, and minimizing the artificial effect of dissociation on gene expression is of critical importance. Here, we compared two proteases, trypsin and CAP, and found that trypsin treatment at 25 °C upregulated Notch target genes regardless of cell type, but CAP treatment did not. This result suggests that trypsin treatment of *Drosophila* gastrula cells induces Notch signal activation by unknown mechanisms. One possible mechanism is the loss of cis-inhibition of Notch (Del Álamo et al., 2011). The other is the direct activation of Notch by trypsin. Since artificial Notch activation by trypsin has also been reported in mammalian cells (Liu et al., 2014), this may be a universal phenomenon in animal cells. Although the detailed mechanism of Notch activation by trypsin is still unclear, these results indicate that cell dissociation methods need to be carefully considered not only for mammalian tissues as previously reported (Adam et al., 2017; Van Den Brink et al., 2017; Denisenko et al., 2020; Miyawaki-Kuwakado et al., 2021; O’Flanagan et al., 2019), but also for insect tissues. We also found that not all Notch target genes were upregulated by trypsin treatment. For example, *E(spl)* complex genes, such as *E(spl)m8-HLH*, were upregulated ubiquitously, but *sim*, another Notch target gene, was not. A possible cause of this difference is that *sim* has a higher threshold and responds more slowly than *E(spl)* complex genes (Falo-Sanjuan et al., 2019). In addition, Notch may be able to instruct the expression of *E(spl)* complex genes in all cells, but *sim* expression is only permitted by Notch and may require other activators (Bray and Furriols, 2001).

### The whole picture of transmembrane-related DEGs between pair-rule stripes

In this study, we succeeded in reconstructing the transcriptome of stripe patterns at the single-cell level using scRNA-seq data, allowing us to seek quantitative differences in the gene expression neighboring cells for all genes. We then determined that each adjacent pair showed 9–25 DEGs. These DEGs were mainly composed of transmembrane genes, in addition to pair-rule TF genes. Since genes encoding cytoplasmic proteins were hardly detected, the trunk ectoderm could deform based solely on differential expression of plasma membrane genes. The transcriptome features of the trunk ectoderm differ more significantly from those of the mesoderm or endoderm, so that the feature may give the trunk ectoderm the characteristic competence to undergo cell intercalation behavior.

Transmembrane-related DEGs between stripes include three Toll receptor genes (*18w*, *Toll-6,* and *Tollo*), *trn*, and *5-HT2A*, which are known to play a role in the regulation of germband extension (Colas et al., 1999; Paré et al., 2014, 2019; Schaerlinger et al., 2007). Our scRNA-seq data provide an opportunity for a quantitative understanding of their functions, rather than qualitative and binarized data. In addition, DEGs include factors that have not been recognized as regulators of germband extension, but are known to be involved in axon guidance. For example, *comm* and *comm2* are involved in axon guidance across the *Drosophila* midline (Keleman et al., 2002, 2005). *Sema5c* is a member of the semaphorins that was originally identified as axonal growth cone guidance molecules (Yazdani and Terman, 2006). There is growing evidence that many transmembrane proteins identified as factors for neural network formation are also involved in epithelial morphogenesis and homeostasis (Cammarota et al., 2020; Hinck, 2004; Vaughen and Igaki, 2016; Yoo et al., 2016). Furthermore, *Sema5c* was recently reported to be involved in the morphogenesis of follicle epithelia (Stedden et al., 2019). In this analysis, only a small number of genes were identified as DEGs. Although it remains possible that some important genes are being missed because the threshold is too high, the limited transmembrane DEGs are expected to be sufficient to organize the dynamics of epithelial morphogenesis in a redundant or cooperative manner.

### Transient intermediate/hybrid state during cell differentiation

Detailed subclustering revealed two potential intermediate-state cells in *Drosophila* gastrulae. One is the cells belonging to “Anterior midgut wntD”, which express both endodermal and mesodermal markers. The other is the cells belonging to “PS14 ectoderm wg” that are likely to be intermediate between PS14 ectoderm and hindgut. These kinds of intermediate/hybrid (or multilineage priming) states have also been identified in the embryogenesis of other organisms by scRNA-seq analysis (Briggs et al., 2018; Farrell et al., 2018; Packer et al., 2019), suggesting that the transient intermediate state is a common step during cell differentiation. It is thought that such intermediate states do not persist, and the cells should eventually differentiate into one of the two states, but how the direction of differentiation is determined is still not well understood. During development, cell differentiation proceeds in parallel with morphogenesis, and we recently found that morphogenesis can modulate cell differentiation (Kondo and Hayashi, 2019). Since regions of two intermediate states that we identified undergo dynamic changes in tissue shape (mesoderm/endoderm invaginations), it is possible that their cell differentiation paths depend on the completion of invagination of the head mesoderm or hindgut at this time. In the future, it is essential to investigate the lineages of these cells in detail using time-series analyses of cell differentiation and morphogenesis.

### Cell fate transformation in *bcd* mutants at the transcriptome level

How a limited number of TFs and signaling generate various cell types during development is still a fundamental biological question. It has been proposed that sequential logic can overcome the bottleneck of combinatorial logic (Letsou and Cai, 2016). At least in this theoretical view, there is a limit to the transcriptome pattern that can be established from only the combination state at the time. However, if we take the sequential logic wherein the time ordering of factors informs the final outcome, the diversity of target configurations dramatically increases even with the same regulatory network. Although the anterior part of *bcd* mutants eventually got the posterior combination of gap genes (Staller et al., 2015), the temporal histories of gap gene expression and pair-rule gene expression are slightly different from those of the original posterior region. For example, in *bcd* mutants, a gap gene *hb* mRNA expression is observed in both the anterior and posterior regions, but the posterior one initiates earlier than the anterior one (Staller et al., 2015). In addition, in *bcd* mutants, pair-rule genes *ftz* and *eve* show mutually exclusive expression patterns at the end of cellularization, but they show extensive overlap in earlier stages. This overlap was more evident than in the WT.

However, our scRNA-seq analysis of *bcd*-depleted embryos revealed that the anterior part of them acquired transcriptome characteristics for cells in the posterior region (Figure 7), strongly suggesting that the temporal histories of gap genes and pair-rule genes do not significantly affect the formation of transcriptome, and the status at last time just before gastrulation starts (50 min after the onset of nuclear cycle 14) determines the cell fate. Therefore, at least from early zygotic control in the *Drosophila* gastrula, the sequential scheme has less contribution, possibly because of the short duration of the process. This supports the proposed possibility that “subsequent layers serve to transform the positional information, fully available already at the gap gene layer, into an explicit commitment to repeated but discrete cell types” (Petkova et al., 2019). Furthermore, even though there are some noise and sharp discontinuities along the A-P axis, all cells in *bcd* mutants eventually canalize into cell types that are present in wild type at the level of the transcriptome, suggesting that the robust gene regulatory mechanism is operating not only with a handful of marker genes, but also with a multitude of genes across the whole genome.

### Non-linear conversion from spatial information to cell-type specific transcriptome

During the process from patterning to cell differentiation, it is widely considered that gradual positional information is converted into the expression of TFs, and these combinatorial patterns determine cellular lineages in a discrete fashion (Briscoe and Small, 2015; Petkova et al., 2019). However, most studies have been based on a limited number of TFs, and the understanding of the relationship between cell differentiation status and genome-wide gene expression profiles is still limited. In this study, we established the scRNA-seq dataset with 64 subcluster information, and the hierarchical clustering of these subclusters revealed that, in terms of the transcriptome level, rather than the expression profiles of TF genes, those of PM-related genes or other effector genes better represented the cell differentiation status corresponding to the three germ layers. This could indicate that these effector genes directly define cellular characteristics, such as behaviors and functions. Furthermore, it is suggested that when viewed from the perspective of only mRNA expression profiles of all TFs, initial positional information still remains there to some extent. In other words, without any prior functional knowledge about each gene, only the mRNA expression profiles of TF genes were not sufficient to distinguish the three germ layers. These analyses using our scRNA-seq data support the idea of a non-linear combinatorial scheme of transcriptome establishment by TFs from the point of transcriptome-wide view. In addition, it also implies that, in order to understand the regulatory mechanisms of cell differentiation, it is not enough to look at the overall mRNA expression of TFs, but it is important to look at the expression profiles of downstream effector genes and to focus the TFs that have a strong contribution. This may be because a small subset of TFs is the true master controller that sets up the lineages and that these are the ones that drive the expression of the cellular effector genes in a non-equivalent manner. One possibility is that mRNA expression levels do not reflect the activity of each TF, and such master regulators are highly activated by post-transcriptional modifications through cellular signaling to have stronger effects than others. This is consistent with the fact that the expression profiles of PM-related genes were significantly variable among cells in the gastrula. Alternatively, from the diverse states of TF expression, a strong combinatorial action of common master controllers emerges in the gene regulatory network, resulting in the induction of the downstream effector gene profile corresponding to each germ layer. It is also possible that similar expression profiles of effector genes arise from different combinations of TFs.

In this study, by using our scRNA-seq data with Perler or NovoSparc, we reconstructed the spatial transcriptome of *Drosophila* gastrula at single-cell resolution with high accuracy. Our scRNA-seq data with spatial reconstruction can be used as an important reference for the elucidation of regulatory networks, including cell-cell communication. Future integrated analyses of the genome-wide gene-regulatory network and spatial signaling activity with this scRNA-seq data will provide us with more detailed insights into the mechanisms by which the gradual positional information is non-linearly converted into discrete patterns of cell differentiation, and also enable a deeper understanding of the developmental systems that orchestrate tissue morphogenesis and functions.

## Acknowledgements

We would like to thank the Kyoto Stock Center, the Bloomington Drosophila Stock Center for fly stocks, Platform for Advanced Genome Science (PAGS), and NGS Core Facility of the Graduate Schools of Biostudies, Kyoto University for supporting the sequencing analysis; Seishi Ogawa, Masahiro Nakagawa, and Toshiko Sato for supporting the 10x Chromium; Masayo Miki for assistance with the experiments; Tadao Usui and Yu-Chiun Wang for critical comments on the manuscript; members of the Uemura laboratory, Shigehiro Kuraku, and the Laboratory for Phyloinformatics RIKEN BDR for discussion. This work was supported by a Grant-in-Aid for Scientific Research (B) (17KT0021 to T.K.), a Grant-in-Aid for Challenging Exploratory Research (15K14535 to T.K.), a Grant-in-Aid for JSPS Research Fellow (20J23385 to S.S.), a Grant-in-Aid for Scientific Research (16H06279 to PAGS) from the Japan Society for the Promotion of Science (JSPS); JST FOREST Program (JPMJFR204V to T.K.), the Naito Foundation (to T.K.), and the Keihanshin Consortium for Fostering the Next Generation of Global Leaders in Research (K-CONNEX) established by the Building of Consortia for the Development of Human Resources in Science and Technology, MEXT (to T.K.). S.S. was supported by a JSPS Research Fellowship for Young Scientists.

## Author Contributions

T.K. conceived the project. S.S. and T.K. performed the experiments and analyzed the data. S.S., Y.O., and H.N. developed and implemented the computational method. C.T., O.N., and M.K. assisted in establishing the scRNA-seq method. T.U. provided feedback on this study. S.S. and T.K. wrote the manuscript with input from all authors.

## Competing Interests

The authors declare no competing interests.

## Materials and Methods

### Fly strains

All stocks were maintained on standard laboratory food containing corn flour, corn grits, dry yeast, glucose, agar, propionic acid, and butyl p-hydroxybenzoat. The following fly strains were used: *y w* for Set1 C1-HT and Set2 trypsin-10x data and *w*; *R14E10-GAL4[attP2] UAS-mCD8.chRFP (III)* for Set3 CAP-10x data was used as a control. Maternal RNAi knockdown of *bcd* was performed as previously reported (Staller et al., 2015). Briefly, *UAS-bcd RNAi* (HMS00035) females were crossed with *matalpha4-GAL-VP16[67]* and *matalpha4-GAL-VP16[15]* males. The *matalpha4-GAL-VP16[67]*/+; *matalpha4-GAL-VP16[15]*/ *UAS-bcd RNAi* (HMS00035) females hatched from it were crossed with *UAS-bcd RNAi* (HMS00035) males, and embryos obtained from this cross were used as *bcd*-RNAi embryos.

### Preparation and fixation of single-cell suspensions

All equipment, including the forceps, brushes, and nylon mesh, was treated with RNase quiet (Nacalai) and washed well with RNase-free water. Embryos were collected by egg laying for 20–30 min and kept for 90 min at 25 °C. Then, embryos were dechorionated using bleach and washed with RNase-free PBS. The developmental stages of embryos were monitored under a fluorescent stereomicroscope (Nikon SMZ18), and stage 6 to 7 embryos shortly after initiation of gastrulation were picked and transferred into 10 µL of ice-cold homogenization buffer (1x RNase-free PBS, 5% trehalose) in a 1.5 mL microtube (Watson, PROKEEP protein low binding tube). Trehalose was included in the whole dissociation process as a cell protectant (Saxena et al., 2012). After collecting 150– 300 embryos at the bottom of the microtubes, the vitelline membranes were broken by slowly turning the tip of the pipette tip (Axygen, Maxymum Recovery 200 μL Universal Fit Tip with Filter). The disrupted embryos were suspended in 500 µL of ice-cold homogenization buffer and pelleted by centrifugation at 800 rcf for 2 min at 4 °C. After removing the supernatant, the pellet was resuspended in 500 µL of ice-cold homogenization buffer, followed by centrifugation at 800 rcf for 2 min at 4 °C.

For trypsin treatment, the pellet was resuspended in 1x trypsine-EDTA (Sigma, T3924) and kept at 25 °C for 10 min. 500μl of ice-cold stopping buffer (1x PBS, 5% trehalose, 0.375 % BSA (WAKO, 012-23881), 0.1 mg/ml trypsin inhibitor (Sigma, T6522)) was added. After washing with 500μl of ice-cold wash buffer1 (1x PBS, 5% trehalose, 0.375 % BSA (WAKO, 012-23881)) twice, the pellet was resuspended with 200 µL of ice-cold loading buffer (1x PBS, 5% trehalose, 0.5 mg/ml ULTRAPURE BSA (Thermo fisher, AM2616), 1/200 RNasin plus (Promega, N2611)).

For CAP treatment, the pellet after homogenization was resuspended in 500μl CAP solution (5 mg/mL Bacillus licheniformis protease (Sigma P5380), 5% trehalose, in 1x PBS), and kept at 6 °C for 30 min. Then, 500 μL of ice-cold wash buffer2 (1x PBS, 5% trehalose, 0.5 mg/ml ULTRAPURE BSA (Thermo fisher, AM2616)) was added. After washing with wash buffer2 four times, the pellet was resuspended in 200 µL of ice-cold loading buffer.

For either trypsin or CAP treatment, the cells suspended in 200 µL of ice-cold loading buffer were filtered through a cell strainer (FLOWMI Cell Strainers for 1000ul Pipette Tip, 40um Porosity) and fixed with 1 ml of CellCover (Anacyte Laboratories) for 1 h at 25 °C, and then kept at 4 °C overnight. The fixed cells were washed with 500 µL of ice-cold loading buffer and resuspended in 100 µL of ice-cold loading buffer. After counting the density of cells using a hemocytometer, the density was adjusted to approximately 200 or 300 cells/µL for Fluidigm C1-HT, or approximately 300 cells/µL for 10x genomics Chromium.

### Single-cell RNA-seq

scRNA-seq library preparations using Fluidigm C1 with the C1 Single-Cell mRNA Seq HT IFC were performed according to the manufacturer’s protocol with some modifications. Before proceeding to the cell lysis step, all 800 capture sites in the IFC were automatically imaged using an Axio Observer.Z.1 (Zeiss) equipped with an Axiocam 105 color (Zeiss) and an electric stage. One modification was custom primers with inserted 8 base UMI for the reverse transcription reaction. Primer sequences are listed in Supplementary Table S2. We also added the ERCC spike-in mix (Thermo Fisher, 4456740) to the Lysis Mix. Another modification was the concentration of the primers used in the library amplification step. A 10-fold lower concentration of enrichment primer was used. The PCR cycle for library amplification was 12. After quality control and quantification using Bioanalyzer and qPCR, the libraries were sequenced with a NextSeq 500 (Illumina), 75 cycle high-output kit v2 (Read1: 15 cycles, Read2: 69 cycles, Index1: 8 cycles, total 92 cycles).

Library preparations using 10x Chromium with the Chromium Next GEM Single Cell 3ʹ Reagent Kits v3.1, were performed according to the manufacturer’s protocol. The PCR cycles were 11 for cDNA amplification and 11 for library amplification. After quality control and quantification using Bioanalyzer and qPCR, the libraries were sequenced using NextSeq 500 (Read1: 28 cycles, Read2, 56 cycles), NovaSeq 6000 (Illumina) (Read1: 28 or 151 cycles, Read2, 91, 98, or 151 cycles), or HiSeqX (Illumina) (Read1: 151 cycles, Read2, 151 cycles). For the trypsin dataset (Set2), the libraries were sequenced using the NovaSeq 6000. For the CAP dataset, the same library was sequenced three times with NextSeq 500, NovaSeq 6000, and HiSeqX. All reads obtained from the three sequencing times were integrated for analysis.

### Analysis of scRNA-seq data of 10x Chromium

Read1, including UMIs and cell barcords, was trimmed to 28 base lengths using fastx_trimmer (FASTX-toolkit, version 0.0.14, http://hannonlab.cshl.edu/fastx_toolkit). Adapter trimming and quality filtering were performed using fastp (version 0.20.1, for NextSeq 500 data with q 20 --cut_tail −l 28 --max_len1 28 --max_len2 55 --trim_poly_g--trim_poly_x, for 10x NovaSeq data -q 20 --cut_tail −l 28 --max_len1 28 --max_len2 97--trim_poly_g --trim_poly_x, for HiSeqX data with -q 20 --cut_tail −l 28 --max_len1 28 --max_len2 97 --trim_poly_x) (Chen et al., 2018). The trimmed reads were mapped to the genome sequence of *Drosophila melanogaster* (BDGP6.22.98) and UMI-counted using STARsolo (version 2.7.7a)(Kaminow et al., 2021). In this process, since STARsolo (version 2.7.7a) cannot account for multi-gene reads for UMI counting, a modified gtf annotation file, in which genes overlapped in the same direction of the genome were integrated and treated as the same gene (Supplementary Table S3), was used. For cell filtering, the median of the total UMI per cell in the filtered output of STARsolo was calculated, and cells with a total UMI two times higher than the mean value were filtered as potential doublets. Then, cells in which either the number of genes detected, the UMI proportion of ribosomal RNA genes, or the UMI proportion of mitochondrial genome genes were outside the range of an average value ± 2.5×SD were filtered as low-quality cells. Remained UMI-count tables were loaded into Seurat (version 3.2.3)(Stuart et al., 2019) and normalized using the SCTransform function with an option (vars.to.regress = c(“percent.mt”, “percent.rRNA”))(Hafemeister and Satija, 2019). “percent.mt” and “percent.rRNA” were labels of metadata which contain the percentages of transcripts from the mitochondrial genome and nuclear rRNA genes to total detected transcripts in each cell respectively. Dimensional reduction analyses were performed using RunPCA and RunUMAP (dims = 1:30, n.neighbors = 20L) functions, followed by unsupervised graph-based clustering with FindNeighbors and FindClusters functions in Seurat. A cluster showing high expression of ribosomal protein genes was filtered out as low-quality cells. Each cluster was manually annotated based on the marker genes identified by the FindAllMarkers function. For subclustering of each cluster, unsupervised graph-based clustering with FindNeighbors and FindClusters functions were further applied to each cluster, as listed in Supplementary Table S1. In general, highly variable features identified using SCTransform were used for clustering. One exception is that subclustering along the DV axis of the lateral ectoderm was performed using k-means clustering (n=7) with only 35 DV genes listed in Supplementary Table S4. Each subcluster was manually annotated based on the marker genes shown in Figrues 2, S3, and S4. During the process of subclustering, cells showing expression of both ectodermal and mesodermal genes were removed as doublets. The remaining singlet dataset consisted of 6,118 cells. All plots were generated using Seurat or ggplot2 in R unless otherwise noted.

### Analysis of scRNA-seq data of C1-HT

Adapter trimming and quality filtering were performed using fastp (version 0.20.1) with options (-q 20 --cut_tail −l 14 --max_len1 14 --max_len2 68 --trim_poly_g --trim_poly_x), and the trimmed reads were mapped to the genome sequence of *Drosophila melanogaster* (BDGP6.22.98) with a modified gtf annotation file as described above and UMI-counted using STARsolo. Only the data derived from the cells determined to be a singlet from the image of the capture site were loaded into Seurat. For each batch, cells in which either the number of genes detected, the UMI proportion of ribosomal RNA genes, the UMI proportion of mitochondrial genome genes, or the UMI proportion of ERCC spike-ins were outside the range of an average value ± 2.5×SD were filtered as low-quality cells. Then, all four batches were integrated and normalized using SCTransform with an option (vars.to.regress = c(“percent.mt”, “percent.rRNA”, “percent.ERCC”)). “percent.ERCC” were labels of metadata which contain the percentage of ERCC to total detected transcripts in each cell. Dimensional reduction, graph-based clustering, and cluster annotation were performed in the same way as the 10x data. All plots were generated using Seurat or ggplot2 (version 3.3.3)(Wickham, 2009) in R, unless otherwise noted.

### Gene Ontology enrichment analysis

For GO enrichment analysis of all high-quality cells, cells in the pole cell cluster were removed and the list of the top 1,500 highly variable features was extracted using Seurat. The gene list was analyzed using g:Profiler (https://biit.cs.ut.ee/gprofiler) with g:SCS algorithm(Raudvere et al., 2019), and significantly enriched terms in cellular components were identified at an FDR threshold of 0.01, and term_size lower than 4,000. For the analysis of highly expressed genes in the *tsr*-high cluster of set2 10x trypsin dataset, 328 highly expressed genes in cluster 10 (Mesoderm_tsr-high) compared to cluster1 (Trunk_mesoderm) were identified using the FindMarkers function with MAST (Finak et al., 2015) at an FDR threshold of 0.01 and logFC threshold of 0.25. The gene list was analyzed using g:Profiler, and significantly enriched terms of the Kyoto Encyclopedia of Genes and Genomes (KEGG) were identified at an FDR threshold of 0.01.

### Clustering analysis with GLAD

The gene list of each GLAD category was downloaded from https://www.flyrnai.org/tools/glad/web/. Since some genes that are listed in the “Transcription factor/DNA binding” category are also listed in other categories, a modified database was prepared to exclude these duplications. The modified GLAD list is presented in Supplementary Table S5. Integrated gene list of “Trans-membrane proteins,” “Receptors,” “Secreted proteins,” and “Matrisome” categories was used as the list of plasma membrane-related genes. Genes listed in GLAD categories other than the “Transcription factor/DNA binding” category and plasma membrane-related genes were considered as other cytoplasmic genes.

For the analysis using only a specific gene set, only the gene of interest from the top 1,500 highly variable features was extracted, and the dimensionality reduction analysis was performed using Seurat, as described above. The cell identity in the UMAP plot was colored using pre-annotated information.

For the hierarchical clustering analysis, the average normalized expression value of each gene for each 64 subcluster was calculated using the AverageExpression function of Seurat. Euclidean distances for all pairs of clusters in log-transformed gene-expression space were calculated using the dist function of R. Then, hierarchical clustering based on the Euclidean distances was performed with the hclust function with the average method. Euclidian distances and the structure of hierarchical clustering were drawn as a heatmap using the heatmap.2 function in the gplot package (version 3.1.1, https://CRAN.R-project.org/package=gplots) of R.

### Assignment of stripe identities of ectoderms along the A-P axis

scRNA-seq data annotated as “trunk ectoderms 2” (see Supplementary Table S1 for details) were extracted. To infer the stripe positions in parasegement, they were analyzed by k-means clustering (n=4) with nine landmark genes of the stripe position (Figure 5A). Then, data in each stripe were divided into odd or even parasegment by k-means clustering (n=2) with *trn* for stripes 1 and 2 (or *pxb* for stripes 3 and 4) and genes positively and negatively correlated with it. K-means clustering was performed with the k-means function in the ClusterR package (version 1.2.2, https://CRAN.R-project.org/package=ClusterR) of R, and the correlation coefficient between all pairs of 1,000 HVGs was calculated with the correlate function in the corrr package (version 0.4.3) of R. All plots were generated by Seurat or ggplot2 (version 3.3.3) in R. DEGs between each adjacent boundary or super boundary were identified using the FindMarkers function with MAST at an FDR threshold of 0.01, and FC threshold of 1.5.

### Filtering of *tsr*-high cells from trypsin-10x data

To remove *tsr*-high cells from Set2 trypsin-10x data and *bcd*-RNAi data, the correlation coefficients between all pairs of 2,000 HVGs were calculated with the correlate function in the corrr package (version 0.4.3) of R. Then, genes positively and negatively correlated with *tsr* were extracted to perform principal component analysis (PCA) and model-based clustering by the Mclust function in mclust package of R (version 5.4.7, https://cran.r-project.org/package=mclust) with options (pca=30, G=2, modelNames=“VVV”). The cluster with high *tsr* expression was filtered out as stressed cells for further data integration.

### Integration of scRNA-seq data

Reference-based integration between control (10x trypsin set2) and *bcd* RNAi scRNA-seq data was performed with the “FindIntegrationAnchors” and “IntegrateData” functions in the Seurat package. Control data were used as a reference. Further dimensional reductions and clustering analysis of the integrated data were performed using the standard procedures of Seurat. Each cluster was manually annotated using the marker gene information identified by the FindAllMarkers function, and the ratio of each cluster was calculated as the ratio of the total number of cells to the number of cells in each cluster.

### Preprocessing scRNA-seq data for spatial reconstruction of gene expression

For Set3 data, because the BDTNP FISH data does not contain pole cells, 123 cells in the “pole_cells” cluster were removed from the dataset. The UMI-count table of the remaining 5,995 cells was renormalized using SCTransform as described above. Then, dimensional reduction analysis was performed using the RunPCA function of Seurat with default settings.

The raw count table (dge_raw.txt) was obtained from *Drosophila* Virtual Expression eXplorer (https://shiny.mdc-berlin.de/DVEX/). This count table was loaded into Seurat and normalized using the SCTransform function without options. Note that, in this data, pole cells were already removed and the UMI counts for mitochondrial and rRNA genes were not included.

For both datasets, a log-scaled count (“data” slot in “SCT” assay of the Seurat object) and HVGs detected by SCTransform were used for spatial reconstruction.

### Selection of ISH reference landmark genes for spatial reconstruction

ISH spatial references were constructed mainly based on the BDTNP database (D_mel_wt atlas_r2.vpc from http://bdtnp.lbl.gov) and DVEX (bdtnp.txt). The DVEX reference was forked from the BDTNP reference, but three genes (*bowl*, *ems*, and *exex*) were only in the DVEX reference.

Both scRNA-seq data were derived from stage 6-7 embryos, while ISH reference data were established for stage 5 embryos. Some genes in the ISH data dynamically changed the expression pattern from stage 5 to stage 6-7. Therefore, genes whose expression patterns significantly changed between the two time points and that could worsen the reconstruction were removed from the reference. As a result, 67 genes remained as landmarks for spatial reconstruction (Supplementary Table S6). In addition, among 3,039 cells in the DVEX reference, eight cells with y < 0 were removed.

### Spatial reconstruction of gene expression by Perler

Perler (version 0.1.0) Python package was obtained from https://github.com/yasokochi/Perler. For both Set3 and NK-data, log-scaled counts and reference above were loaded into the PERLER object, and then the EM algorithm was performed using the em_algorithm method with option (optimize_pi = False). Next, the distances between the scRNA-seq data points and reference data points were calculated using the calc_dist method with default parameters. Optimization of hyperparameters was performed by the loocv method with default parameters and the gridsearch method with parameters (grids = ((0,1), (0.01,1))). Finally, spatial gene expression patterns were reconstructed by the spatial reconstruction method with parameters (mirror = False, _3d = True, z_scored = False).

### Spatial reconstruction of gene expression by NovoSpaRc

NovoSpaRc (version 0.4.3) reconstruction was mainly performed according to https://github.com/rajewsky-lab/novosparc. First, log-scaled counts and reference were loaded into the Tissue object of NovoSpaRc. Cost matrices for the optimal transport framework were calculated by the set_up_smooth_costs method based on 30 principal components (PCs) and the setup_linear_cost method with the reference and default parameters. Then, spatial reconstruction was performed using the reconstruction method with parameters (alpha_linear=0.3, epsilon=5e-3). Note that the alpha_linear parameter and the number of used PCs were determined by grid search so that the LOOCV score described below was maximized.

### Leave-one-gene-out cross-validation (LOOCV)

Each of the 67 landmark genes in the ISH reference was removed from the reference as the true expression, and spatial reconstruction by Perler or NovoSpaRc was performed using the remaining 66 genes as the reference. Pearson correlations between the reconstructed expression pattern of the removed gene and the truth were then calculated.

For decision of hyperparameter for NovoSpaRc, we calculated the score:

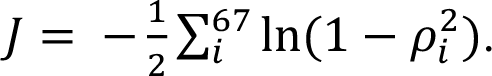

Here, *ρ_i_* is the Pearson correlation coefficient between the ISH expression and the reconstructed expression of gene *i*. The hyperparameter and number of PCs with the highest score were selected.

### Comparison of gene-gene correlation structure conservation

First, common 372 HVGs included in the top 500 HVGs of both Set3 and NK-data scRNA-seq datasets were selected. Pearson correlation between the gene and 371 HVGs in the original scRNA-seq data and those in the Set3-based or NK-data-based reconstruction were calculated. Then, the Pearson correlation coefficient between these two correlation scores for each of the 372 HVGs was calculated as gene-gene correlation structure conservation between the original scRNA-seq data and either reconstruction. For the comparison between Perler and NovoSpaRc, the top 500 HVGs of the Set3 data were used.

### Plotting reconstructed expression pattern

Plotting was performed using the scatter function in the matplotlib (version 3.3.4) package (Hunter, 2007). The reconstructed expression values were converted from a log-scale to a linear scale. For the lateral view, all the cells in the reference were plotted. For dorsal and ventral views, cells with z > 0 and z <= 0 were used, and the cells were mirrored on the x-y plane. The anterior is left in all plots, and the dorsal is up in lateral views.

### Density plot of *ind* and *vnd* expression

For the plot of single cell data, intermediate or medial neuroectoderm cells in the abdomen and PS13 and midline cells were extracted from Set3 data. For the reconstruction data plot, cells with |x| < 50 and −55° < θ < 0° were extracted. θ is the angle between the y-axis and the line segment drawn from the center of the embryo to the cell in a cross section parallel to the yz-plane containing the cell, expressing the position of the cell on the DV-axis (Figure S7G). Density estimation was performed using the gaussian_kde class in the stats module in the Scipy package (version 1.6.0)(Virtanen et al., 2020). Estimation results were plotted using the pcolormesh function in the matplotlib package (version 3.3.4).

### Bulk RNA-seq

For total RNA preparation from embryos, 80 stage 6-7 embryos were harvested, and total RNA was purified using RNeasy Lipid Tissue Mini Kit (Qiagen). For total RNA preparation from dissociated cells, 200-300 embryos at stage 6-7 were dissociated into single-cell suspensions by trypsin-EDTA treatment as described above. After washing, the cells were passed through a 40 µm strainer and pelleted. Total RNA was purified from approximately 40,000 cells using the RNeasy Mini Kit (Qiagen). For total RNA preparation from fixed cells, 200-300 embryos at stage 6-7 were dissociated into single-cell suspensions by trypsin-EDTA treatment and fixed by CellCover as described above. Cells were stored at 4 °C for one day and pelleted. Total RNA was purified from approximately 40,000 pelleted cells using an RNeasy Mini Kit (Qiagen).

cDNA was synthesized from 250 ng of each total RNA using the SMART-Seq v4 Ultra Low Input RNA Kit (Clonetech). Then, a library for Illumina sequencers was constructed from 0.0625 ng cDNA using the Illumina Nextera XT DNA Library Preparation Kit. The libraries were sequenced on an Illumina NextSeq 500 to obtain single-end reads with a length of 76 bases. Each sample was analyzed in duplicates. For each library, 36,577,021–41,844,986 reads were sequenced.

### Analysis of bulk RNA-seq data

Sequenced reads were quality trimmed using Trim Galore (version 0.6.4, https://www.bioinformatics.babraham.ac.uk/projects/trim_galore/) and Cutadapt (version 1.18)(Martin, 2011) with the --nextseq 20 option. After removing the 76th base from each read, the remaining reads were mapped to the genome sequence of *Drosophila melanogaster* (BDGP6.22.98) using STAR with a modified gtf annotation file, as described above. Gene expression was calculated using RSEM (version 1.3.3)(Li and Dewey, 2011), and differential expression analysis was performed using edgeR (version 3.32.1)(Robinson et al., 2009). After removing the mitochondrial and ribosomal RNA genes, low expression genes with CPM less than 0.1 in all six samples were also filtered out. Normalization was performed using calcNormFactors. Spearman correlation coefficients were calculated using the cor function in R. DEGs were identified using the glmQLFit and glmQLFTest functions in the edgeR package at an FDR threshold of 0.01 and logFC threshold of 2.

## Data Availability

All raw sequence data that were deposited in the DDBJ Sequence Read Archive (DRA) and the processed data will be available at the time of publication.

---

**Supplementary Figure S1:**
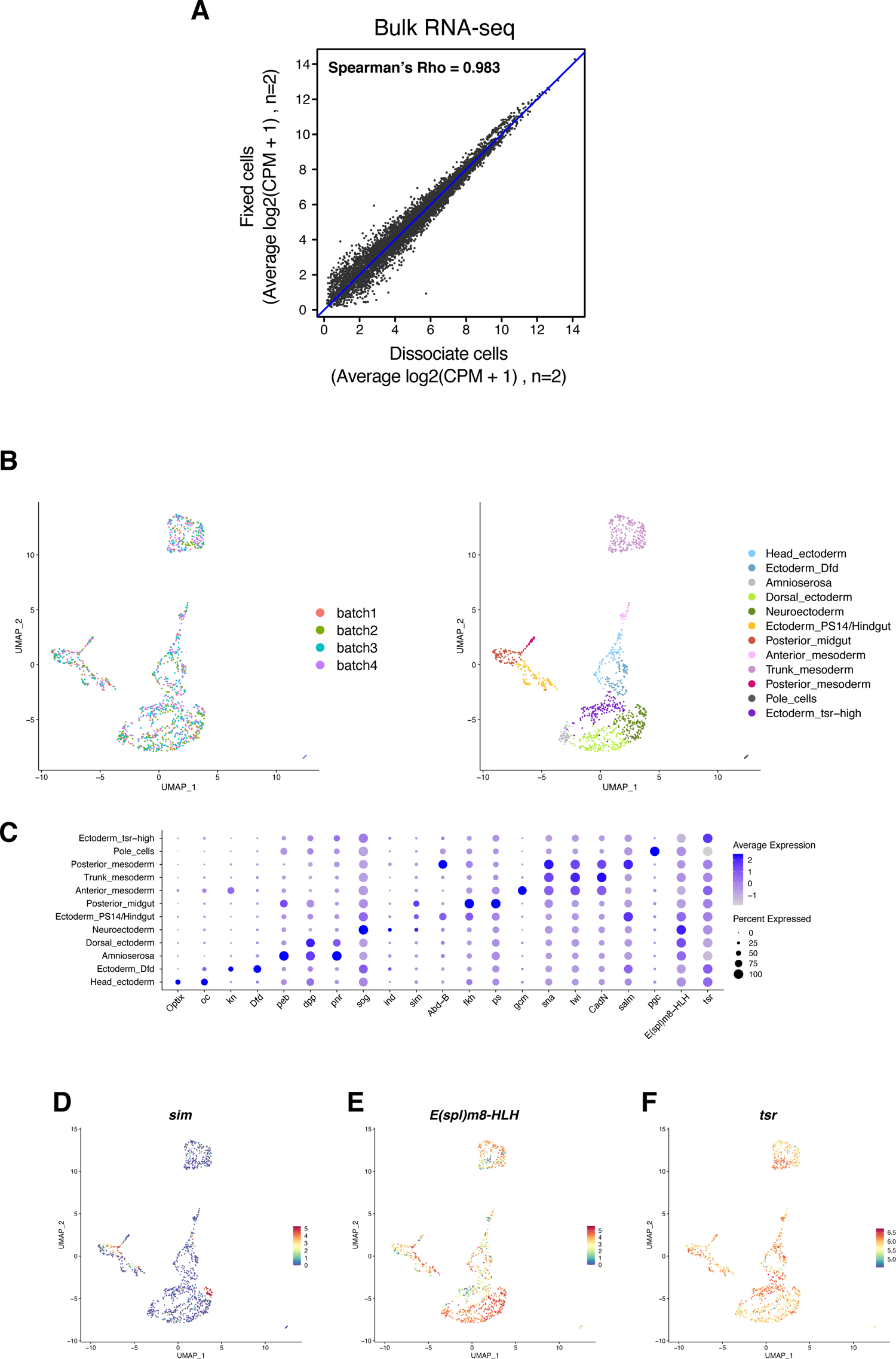
Evaluation of CellCover fixation and C1HT scRNA-seq. A. Scatter plot of bulk RNA-seq data between dissociated-cell and fixed-cell samples. Each dot represents a log2 value of average (CPM + 1). B. UMAP plots of the Set1 trypsin-C1HT scRNA-seq data. Cells were colored by the batch of C1HT run (left) or with cluster information (right). C. Dot plot showing the expression patterns of typical marker genes for each cluster. D-F. Expression patterns of *sim* (D)*, E(spl)m8-HLH* (E), and *tsr* (F) in Set1 trypsin-C1HT data.

**Supplementary Figure S2:**
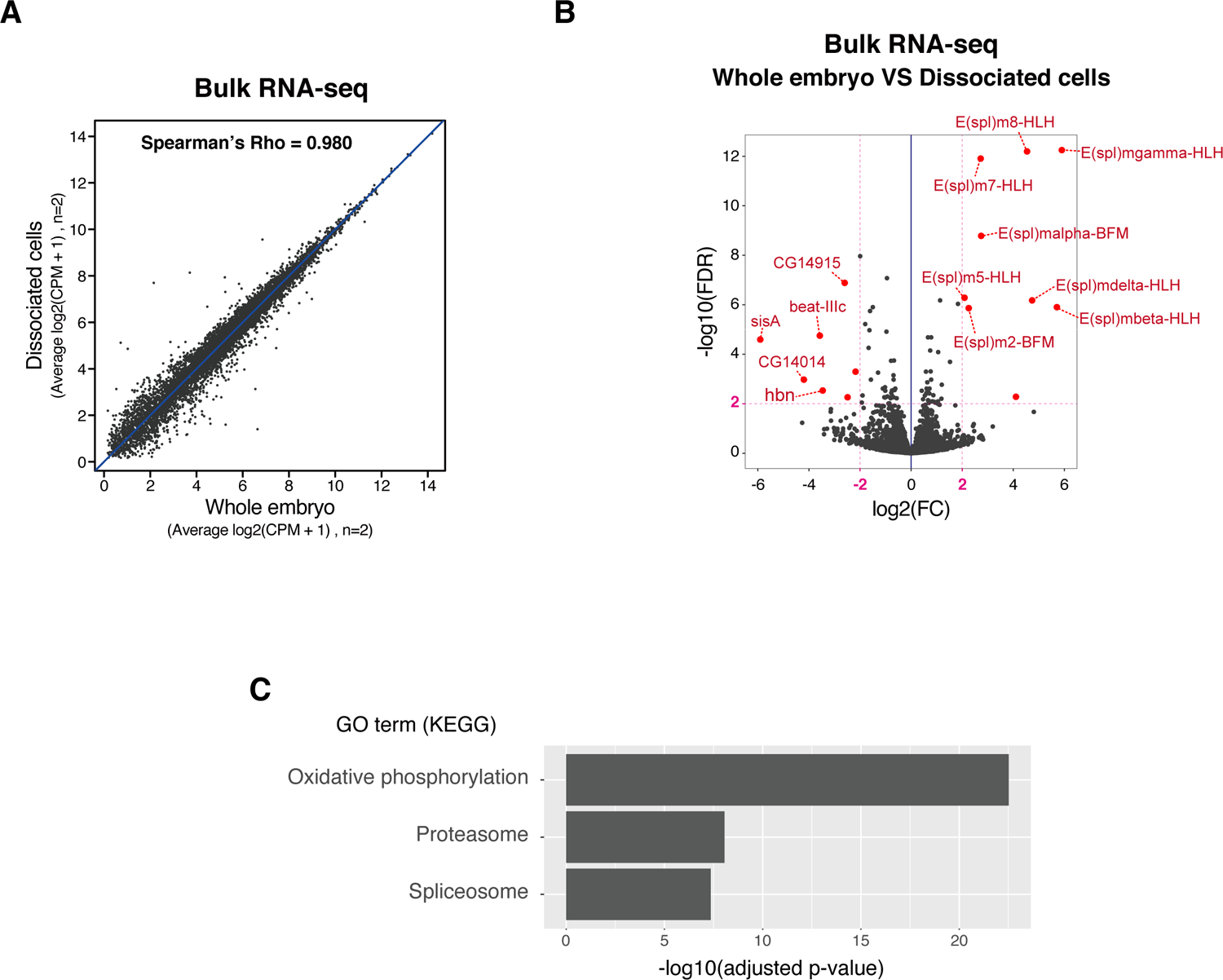
Artificial effect of trypsin treatment. A. Scatter plot of bulk RNA-seq data between embryo and dissociated-cell samples. Each dot represents a log2 value of average (CPM + 1). B. Volcano plot of bulk RNA-seq data between the embryo and dissociated-cell samples. Red dots represent the differentially expressed genes (DEGs) (|logFC| > 2 and FDR < 0.01). C. Gene Ontology enrichment analysis of highly expressed genes in cluster 10 compared to cluster 1 in Set2 trypsin-10x scRNA-seq data. The terms of KEGG are presented.

**Supplementary Figure S3:**
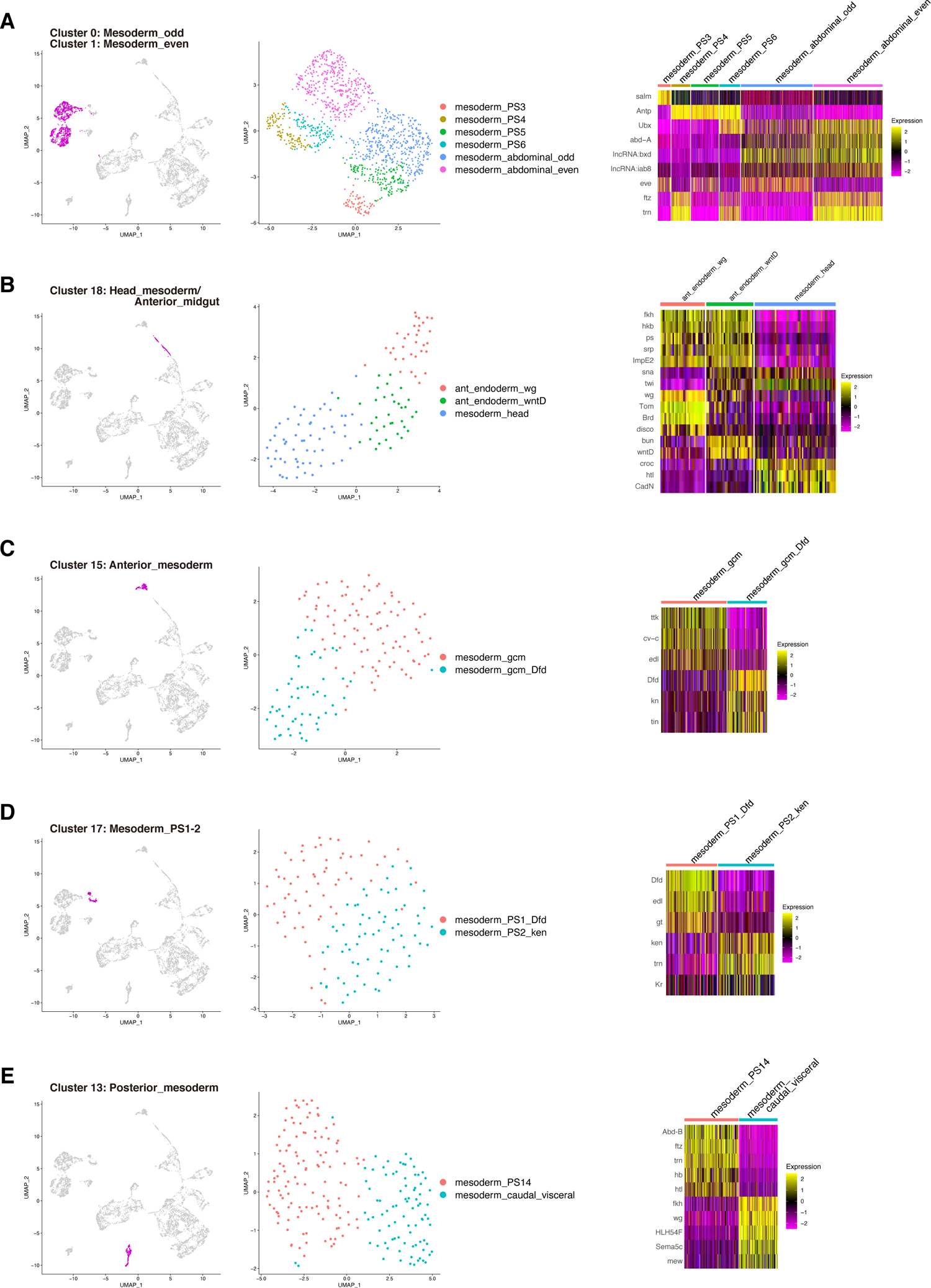
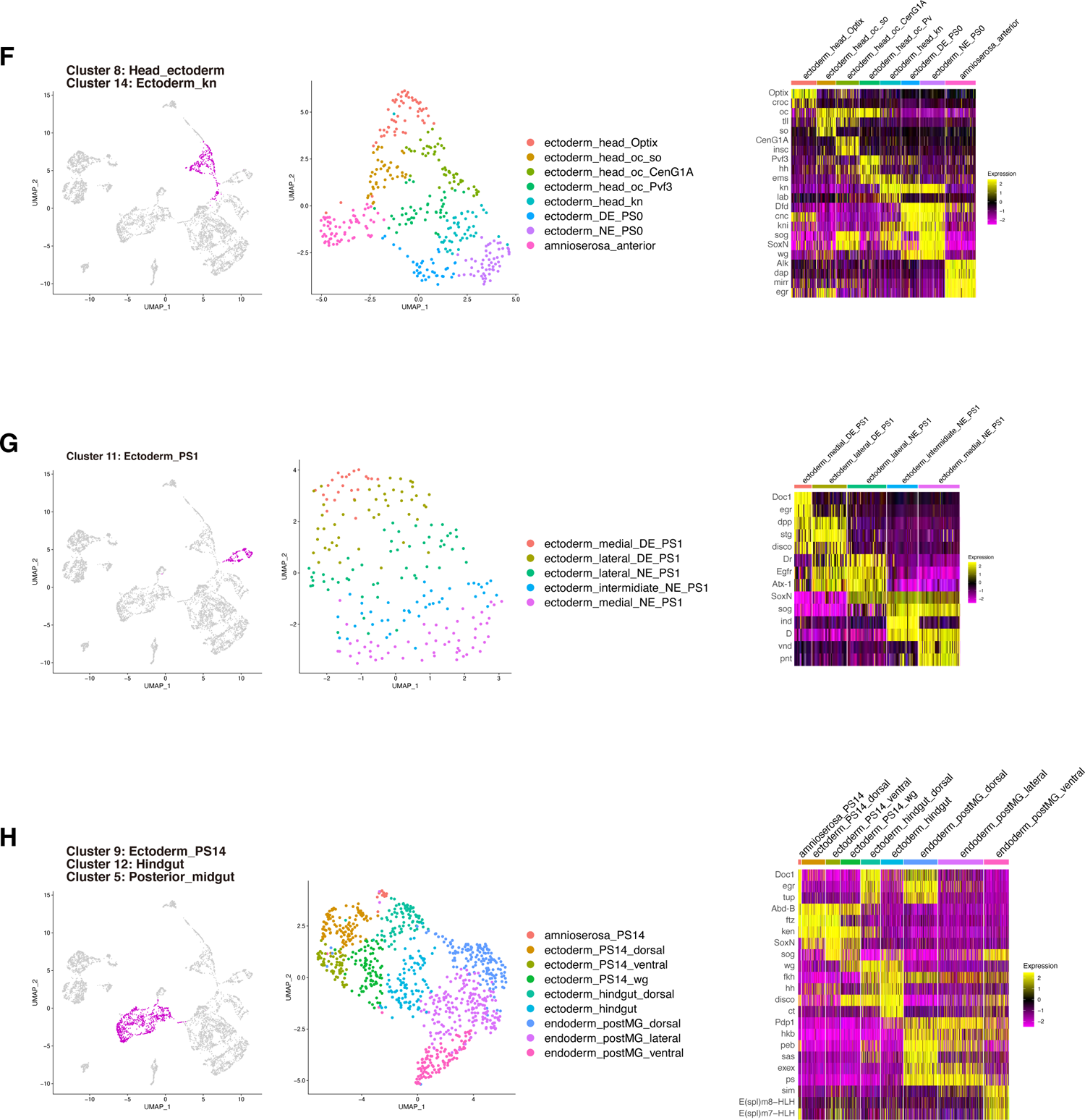
Subclustering of the Set3 CAP-10x data. A. (left) The trunk mesodermal cells from clusters 0 and 1 are colored magenta in the UMAP plot. (middle) UMAP plot and subcluster information for trunk mesodermal cells. (right) Heatmap showing the typical marker genes for all subclusters of the trunk mesodermal cells. B. (left) The head mesodermal/anterior endodermal cells from cluster 18 are colored magenta in the UMAP plot. (middle) UMAP plot and subcluster information for head mesodermal/anterior endodermal cells. (right) Heatmap showing the typical marker genes for all subclusters of the head mesodermal/anterior endodermal cells. C. (left) The *gcm*+ anterior mesodermal cells from cluster 15 are colored magenta in the UMAP plot. (middle) UMAP plot and subcluster information for *gcm*+ anterior mesodermal cells. (right) Heatmap showing the typical marker genes for all subclusters of the *gcm*+ anterior mesodermal cells. D. (left) The PS1 and PS2 mesodermal cells from cluster 17 are colored magenta in the UMAP plot. (middle) UMAP plot and subcluster information for PS1 and PS2 mesodermal cells. (right) Heatmap showing the typical marker genes for all subclusters of the PS1 and PS2 mesodermal cells. E. (left) The posterior mesodermal cells from cluster 13 are colored magenta in the UMAP plot. (middle) UMAP plot and subcluster information for posterior mesodermal cells. (right) Heatmap showing the typical marker genes for all subclusters of the posterior mesodermal cells. F. (left) The head ectodermal cells from clusters 8 and 14 are colored magenta in the UMAP plot. (middle) UMAP plot and subcluster information for head ectodermal cells. (right) Heatmap showing typical marker genes for all subclusters of the head ectodermal cells. G. (left) The PS1 ectodermal cells from cluster 11 are colored magenta in the UMAP plot. (middle) UMAP plot and subcluster information for PS1 ectodermal cells. (right) Heatmap showing typical marker genes for all subclusters of the PS1 ectodermal cells. H. (left) The posterior ectodermal and endodermal cells from cluster 5, 9, and 12 are colored magenta in the UMAP plot. (middle) UMAP plot and subcluster information for posterior ectodermal and endodermal cells. (right) Heatmap showing the typical marker genes for all subclusters of the posterior ectodermal and endodermal cells.

**Supplementary Figure S4:**
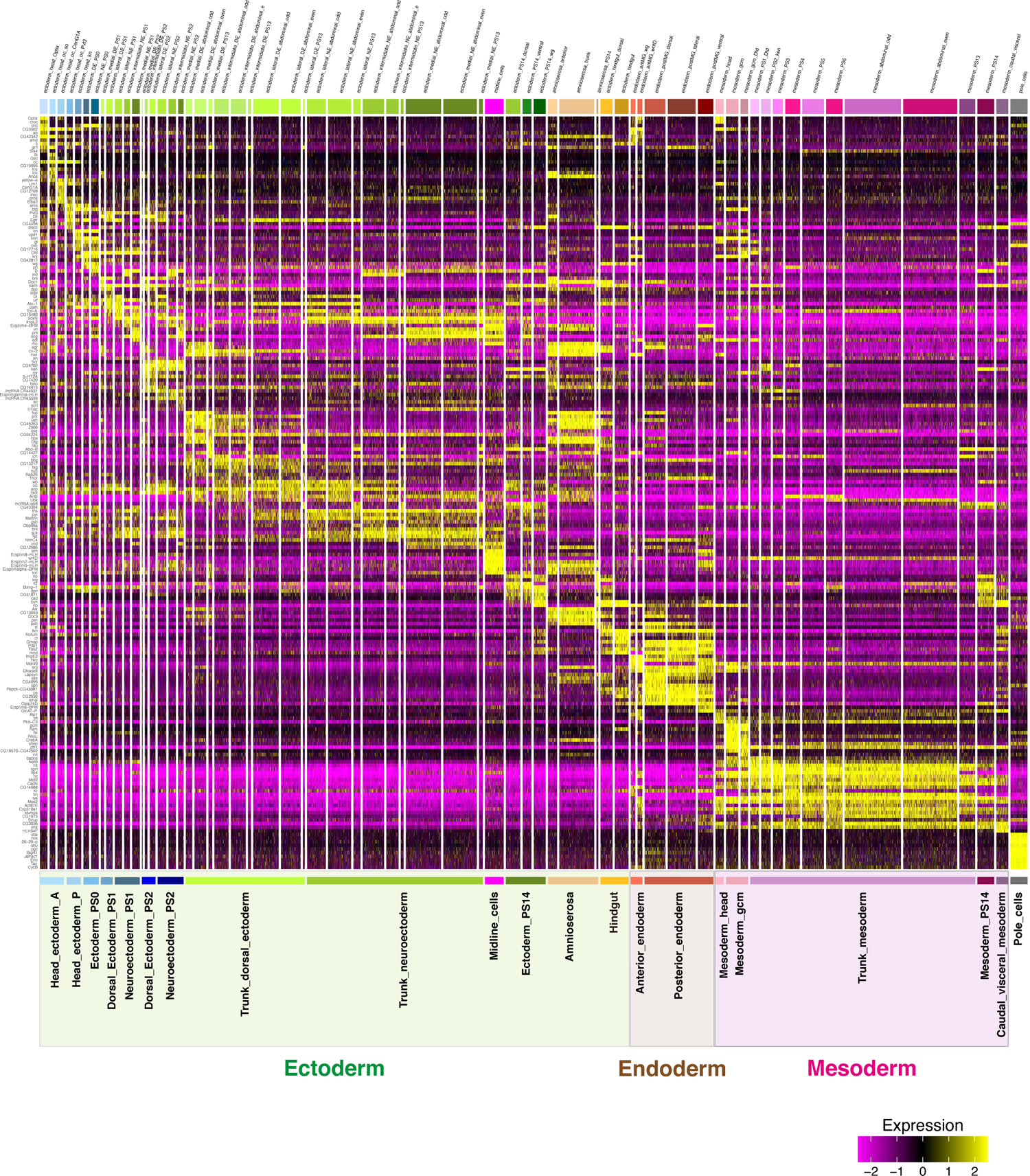
Marker genes for all 65 subclusters. Heatmap showing the top 10 marker genes for all subclusters.

**Supplementary Figure S5:**
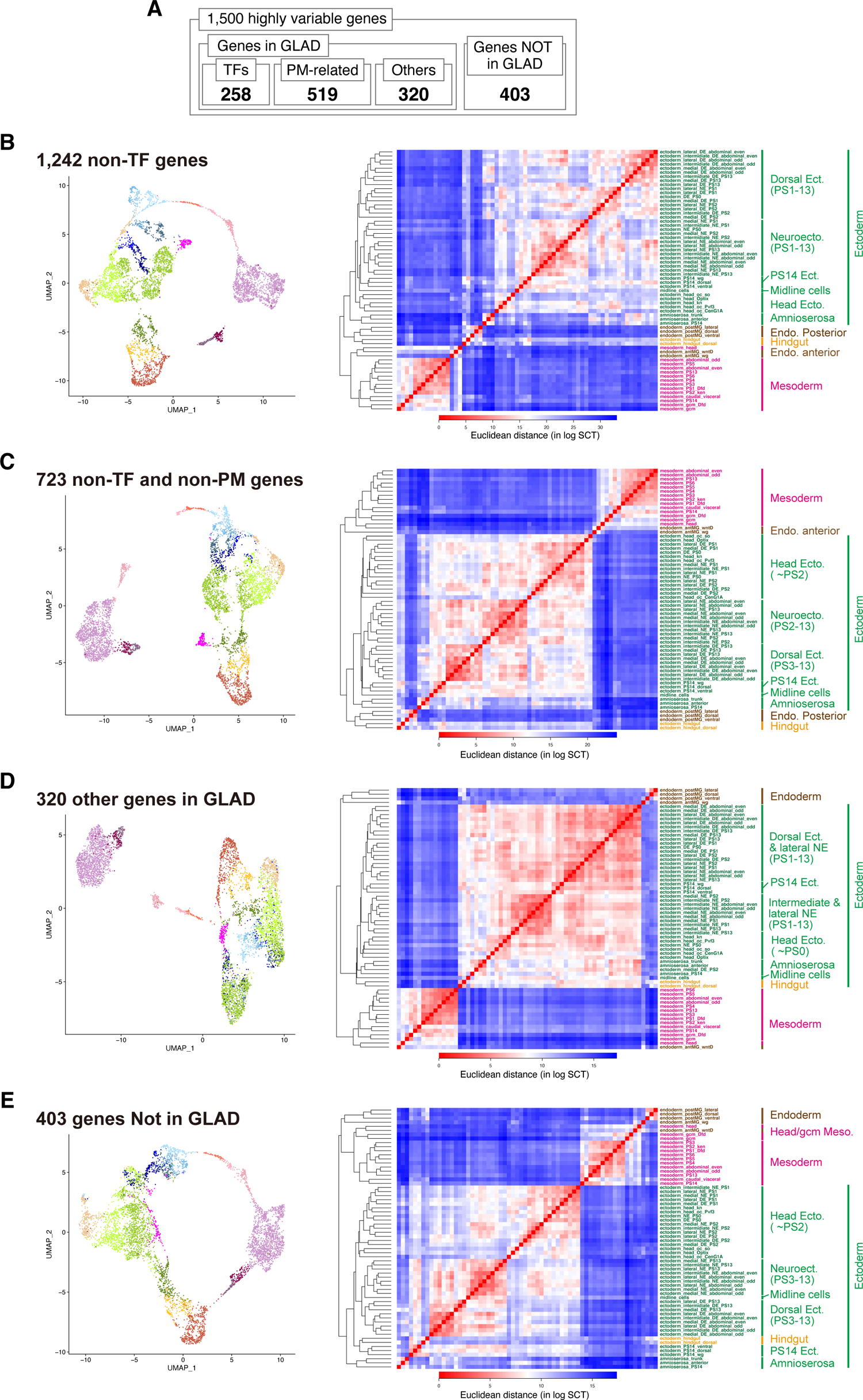
Clustering analysis with GLAD categories. A. Breakdown of GLAD category affiliation for 1,500 highly variable genes. B-E. (left) UMAP plot and with 1,242 non-TF HVGs (B), 723 non-TFs and non-PM HVGs (C), 320 other HVGs in GLAD (D) and 403 HVGs not in GLAD. Cells were colored by super cluster information based on prior annotations (see Figure 4I and Supplementary Table S1 for details). (right) Hierarchical clustering analyses of 64 subclusters based on the Euclidean distances in log-transformed gene-expression space. Subclusters were colored based on the future germ layers.

**Supplementary Figure S6:**
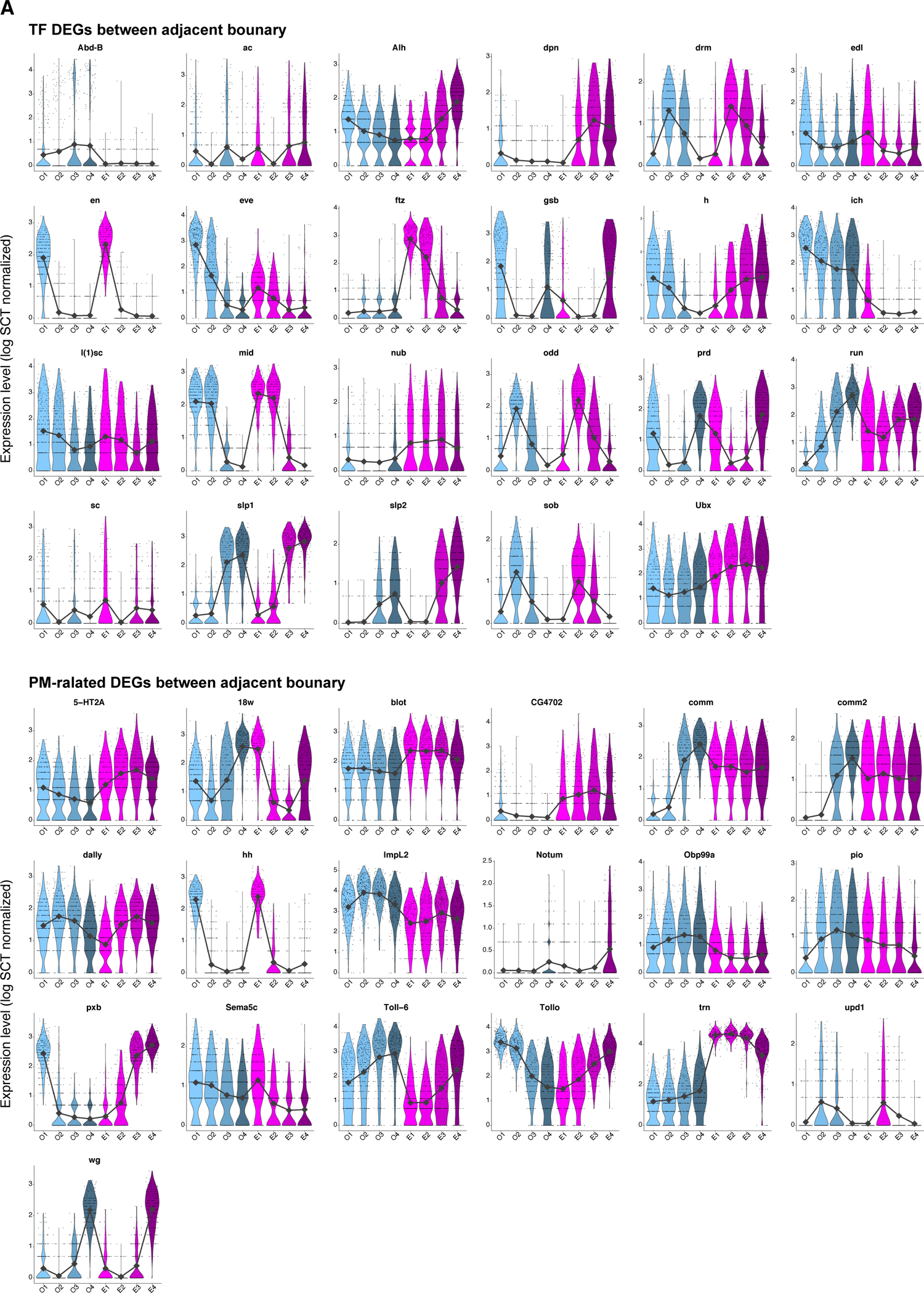
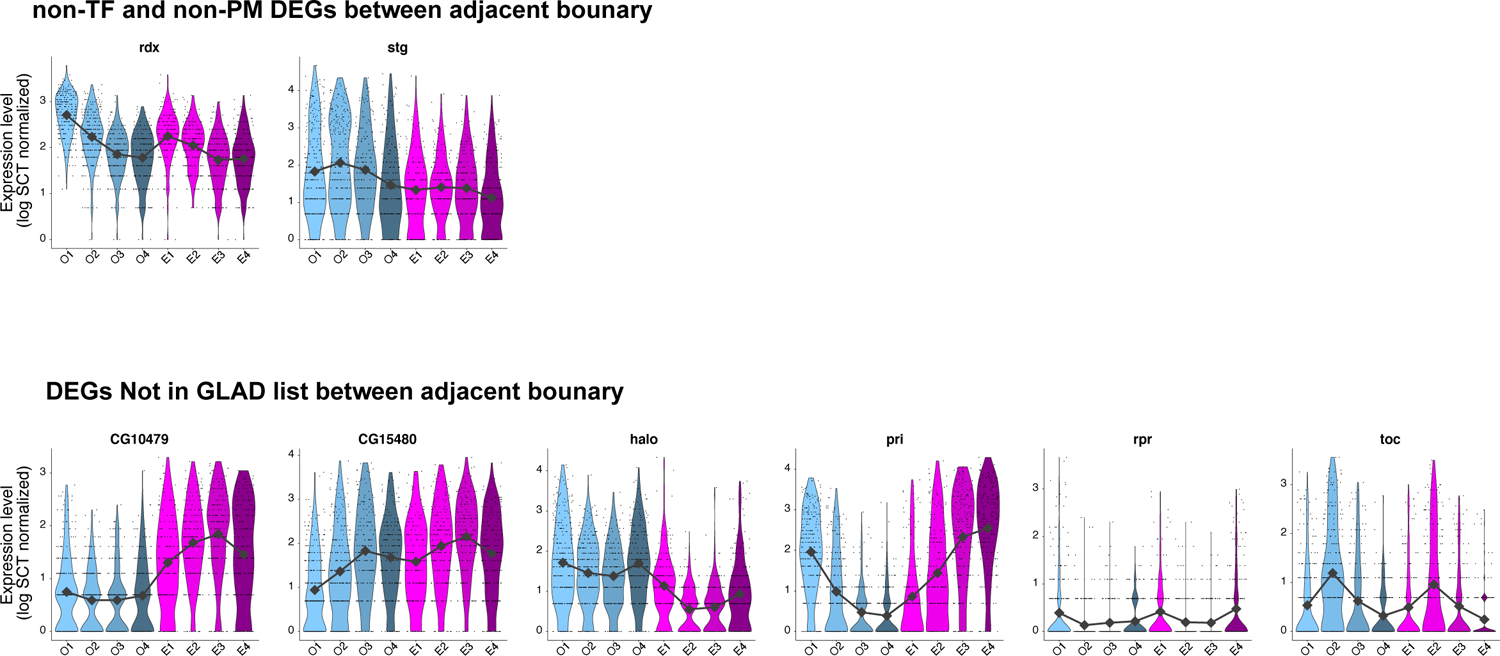
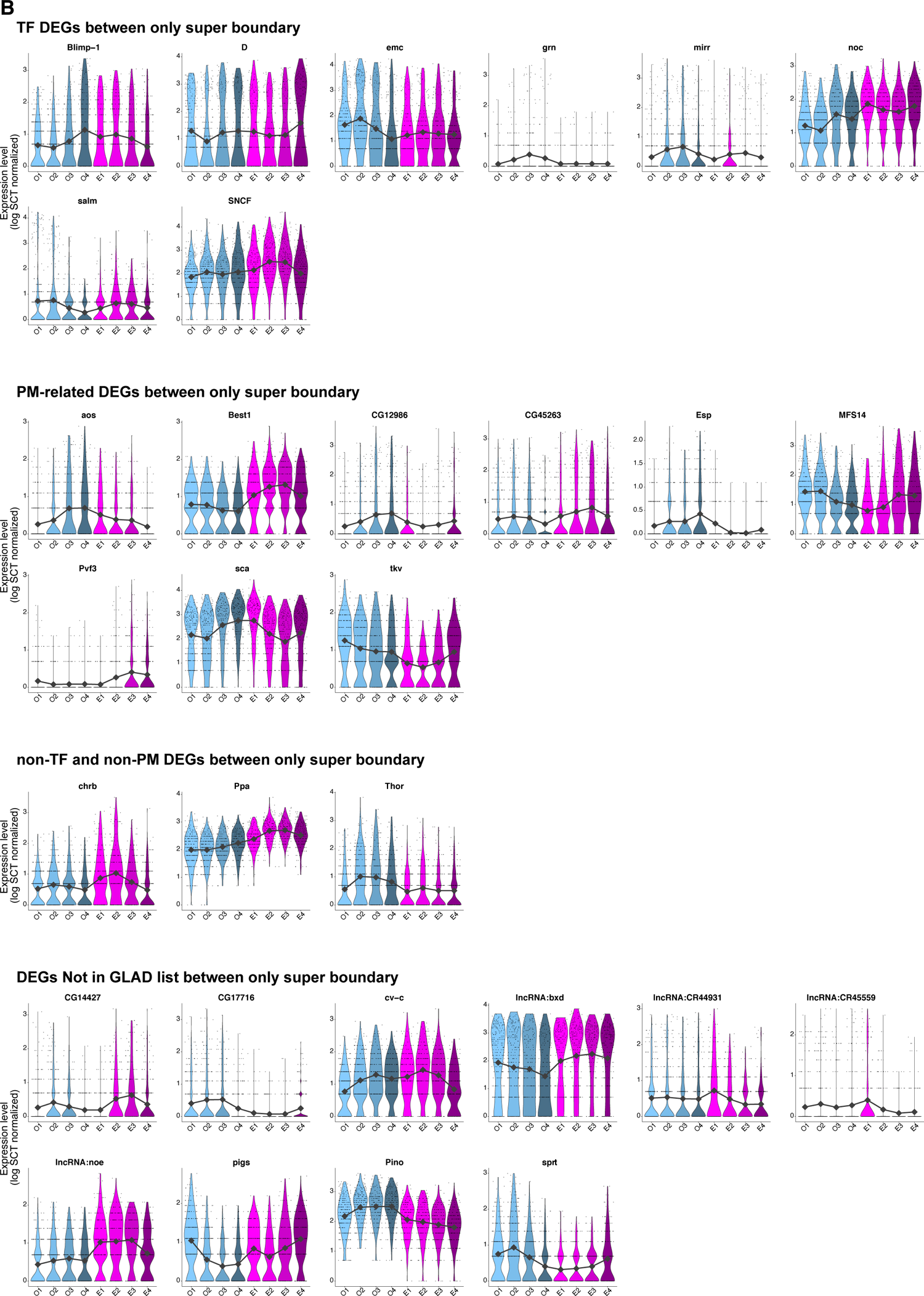
The whole list of DEGs between stripes. A. Violin plots showing the expression patterns of genes identified as DEGs between adjacent boundaries. The gray line indicates the median values for each stripe. Expression levels represent the log-transformed values after SCTransform normalization. B. Violin plots showing the expression patterns of genes identified as DEGs only between super boundaries. The gray line indicates the median values for each stripe. Expression levels represent the log-transformed values after SCTransform normalization.

**Supplementary Figure S7:**
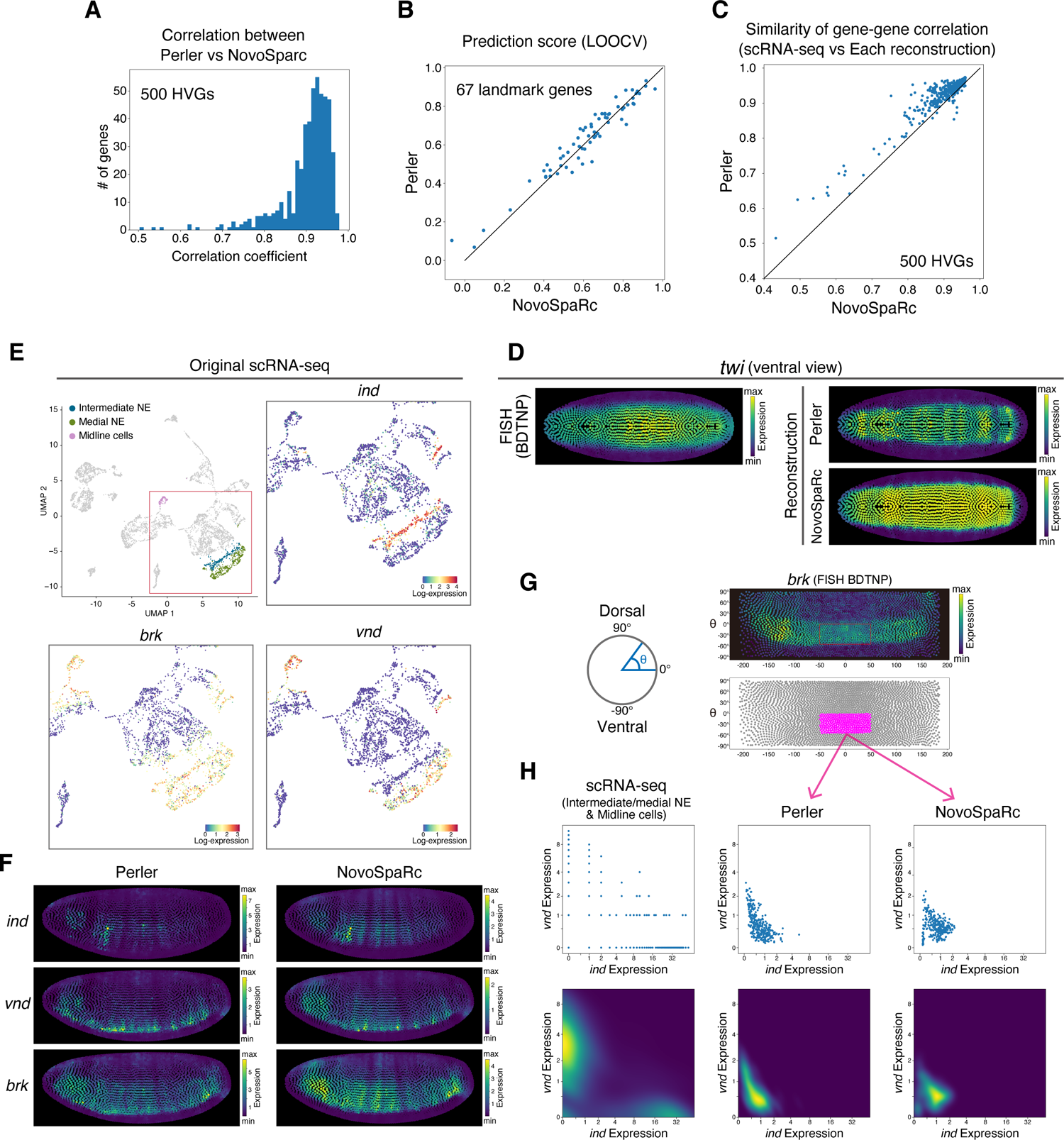
Comparison between the reconstructions by Perler and NovoSpaRc. A. The histogram of Pearson correlation coefficients of each gene expression in top 500 HVGs between Perler and NovoSpaRc reconstruction based on Set3 data. B. Comparison between LOOCV results of reconstructions by NovoSpaRc and those by Perler based on Set3 data. Each dot indicates each gene in reference. X-axis and Y-axis show the Pearson correlation coefficient between the reference expression and reconstruction result of each gene by NovoSpaRc and Perler, respectively. C. Comparison of gene-gene correlation structure conservation between Set3-based reconstruction by Perler and NovoSpaRc. Each dot indicates each gene in top 500 HVGs of Set3 data. The X-axis shows gene-gene correlation structure conservation in the NovoSpaRc reconstruction and the Y-axis shows gene-gene correlation structure conservation in the Perler reconstruction. The definition of gene-gene correlation structure conservation was described in Materials and Methods. D. Reference or reconstructed expression patterns of *twi*. Colormap is linear and min-max scaled. Ventral view. E. Upper-left panel shows the cells used for the plot in (H, left). Other panels show the expression patterns of *ind* (upper-right), *brk* (bottom-left), and *vnd* (bottom-right). F. Reconstructed expression patterns of *ind* (top), *vnd* (middle), and *brk* (bottom) by Perler (left) and NovoSpaRc (right). G. (left) Schematic diagram showing the definition of the position of a cell along the DV axis. (upper-right) Reference expression patterns of *brk*. The embryo is shown in a flattened view. (bottom-right) Magenta indicates cells within the red rectangle region in the upper-right panel. These cells were used for the plot in (H, middle and right). H. (Top) Scatter plots of the expression patterns of *ind* (X-axis) and *vnd* (Y-axis) in each neuroectoderm or midline cell in scRNA-seq (left), Perler reconstruction (middle), and NovoSpaRc (right). (Bottom) Density plots of the expression levels of *ind* (X-axis) and *vnd* (Y-axis) in the neuroectoderm and midline cells in scRNA-seq (left), Perler reconstruction (middle), and NovoSpaRc (right). Color map is min-max scaled. In both plots, each axis is shown in the log-scale and the scale labels are linear values.

**Supplementary Figure S8:**
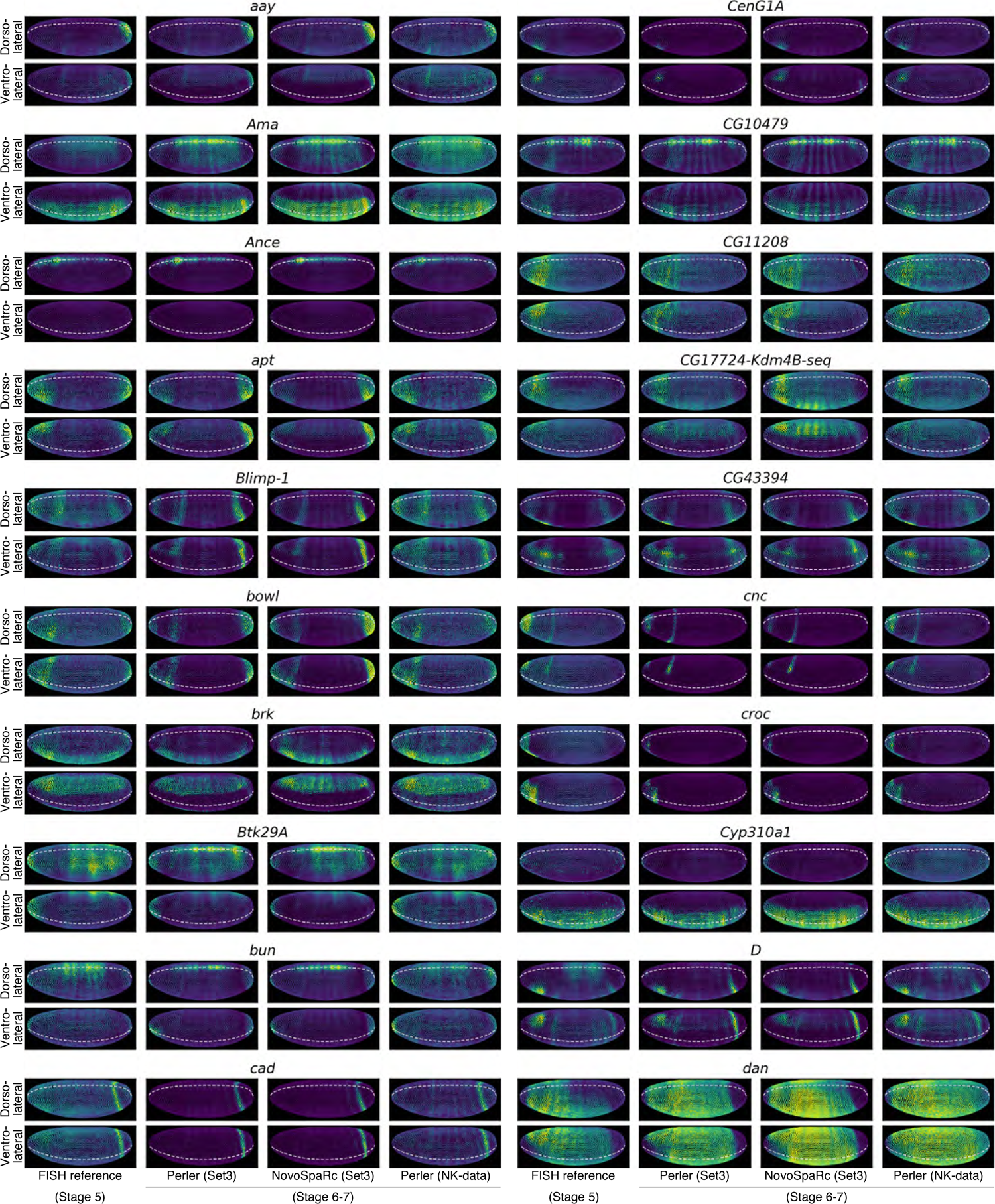
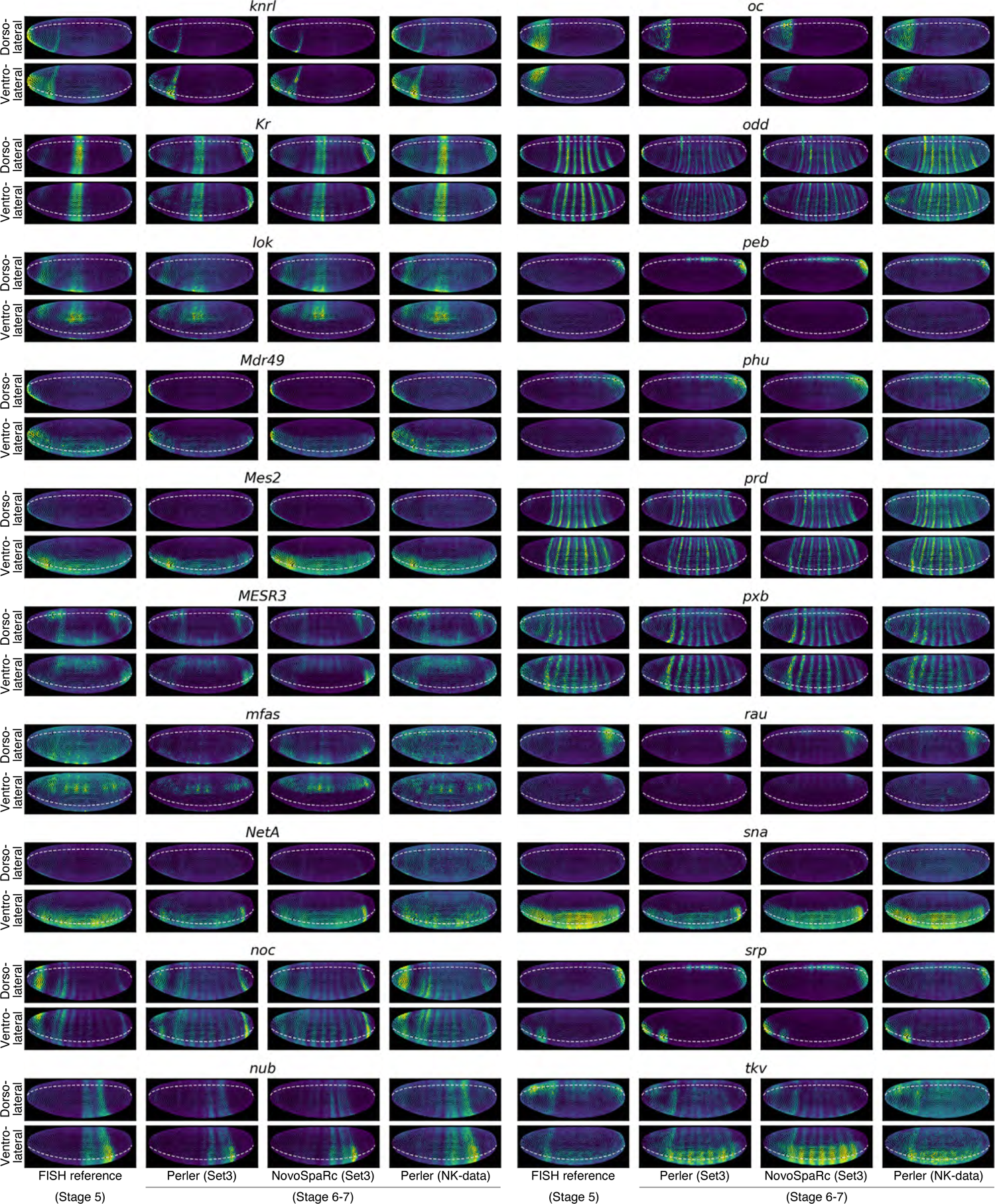
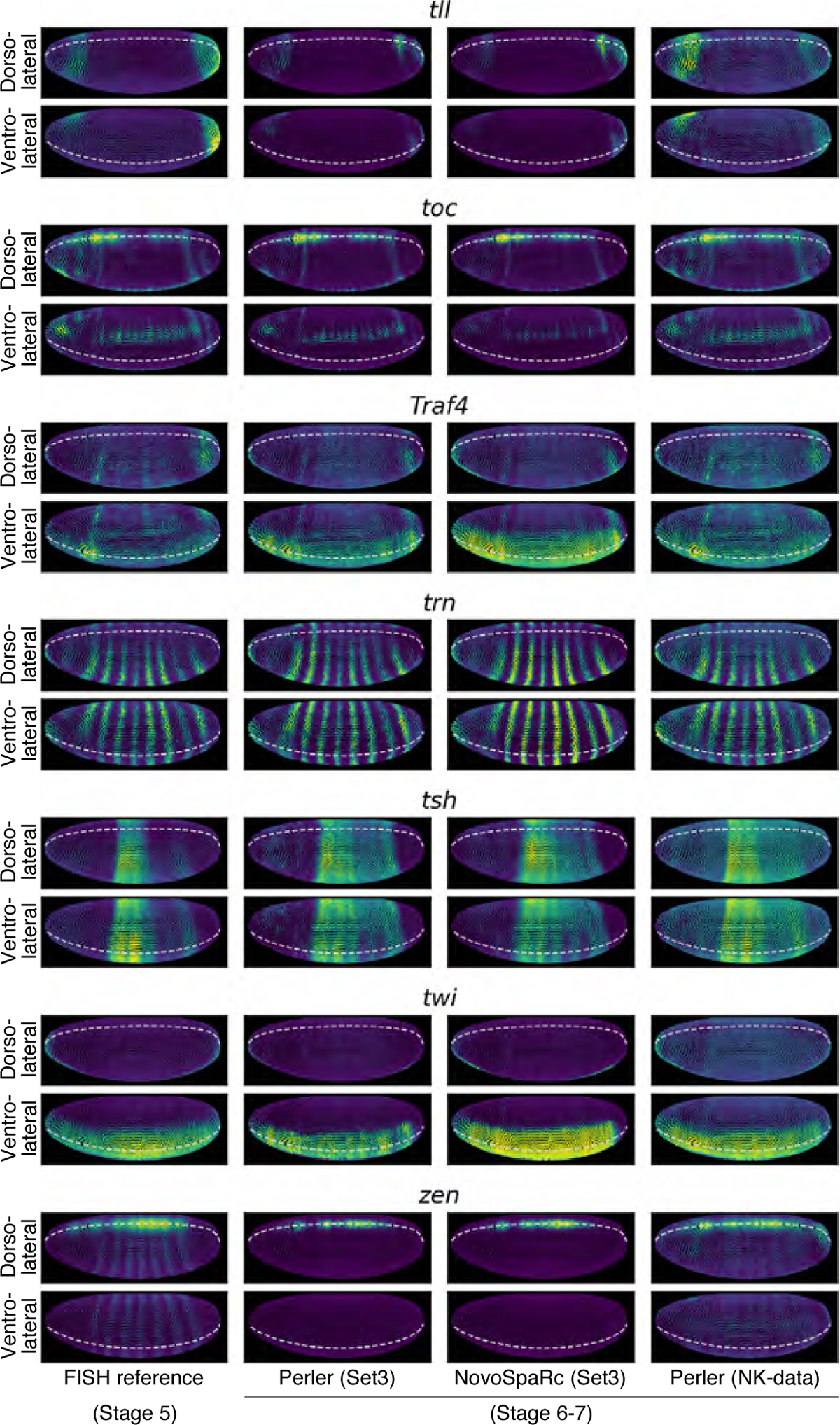
Spatial reconstruction of landmark genes. ISH reference expression patterns (left) and spatial reconstruction of all landmark genes (67 genes) by Perler with Set3 data (middle-left), NovoSpaRc with Set3 data (middle-right), and Perler with NK-data (right). Colormaps are linear and min-max scaled. The upper panels show the dorsolateral views (θ = 45°, Figure S7G), and the lower panels show the ventrolateral views (θ = –45°, Figure S7G). Dashed lines indicate the dorsal or ventral midline.

**Supplementary Table S1.**
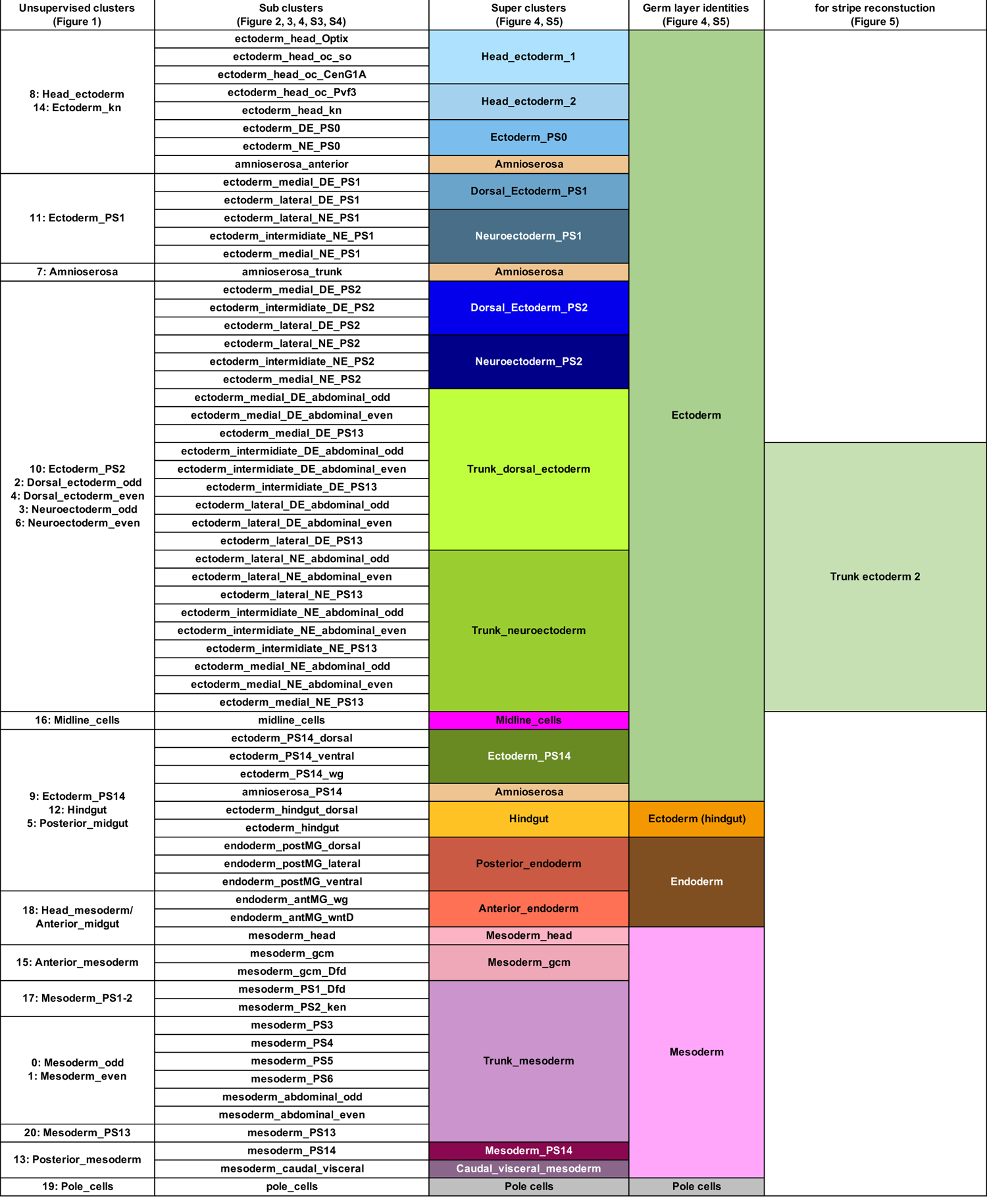

**Supplementary Table S2.**
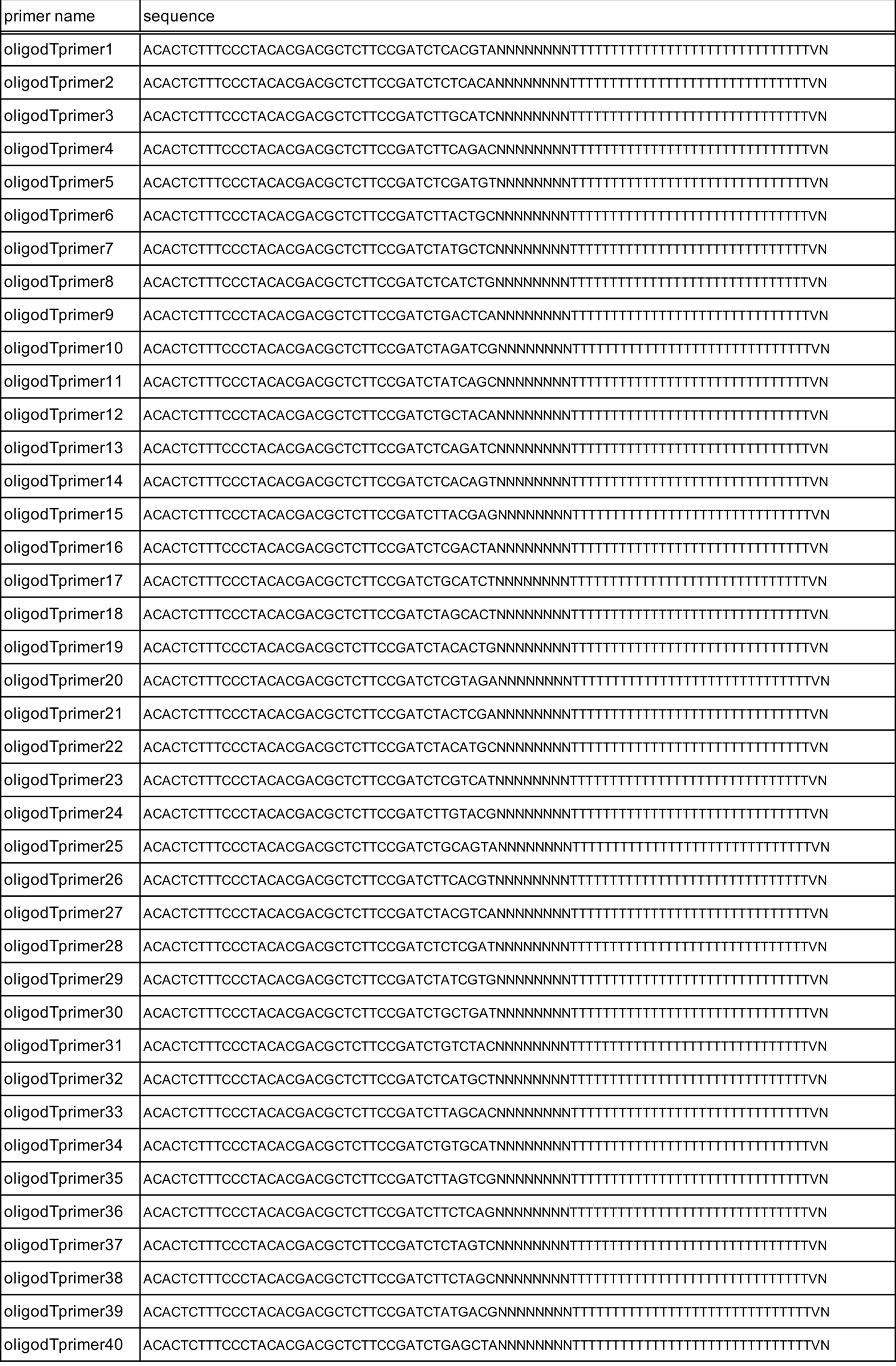

**Supplementary Table S3.**
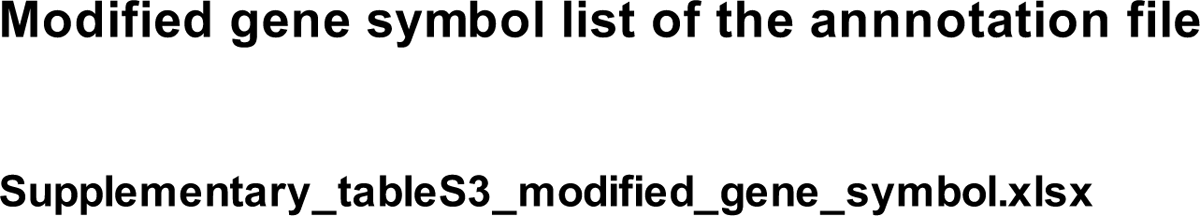

**Supplementary Table S4.**

|  |  |  |  |  |  |  |
| --- | --- | --- | --- | --- | --- | --- |
| *Ama* | *Ance* | *Atx-1* | *bbg* | *brk* | *C15* | *CG13653* |
| *cic* | *cv-2* | *dap* | *Doc1* | *Doc2* | *Doc3* | *dpp* |
| *Dr* | *Dtg* | *Egfr* | *egr* | *emc* | *ind* | *mirr* |
| *peb* | *pnt* | *pnt* | *rho* | *sog* | *SoxN* | *srp* |
| *stg* | *tup* | *ush* | *vn* | *vnd* | *Z600* | *zen* |

**Supplementary Table S5.**
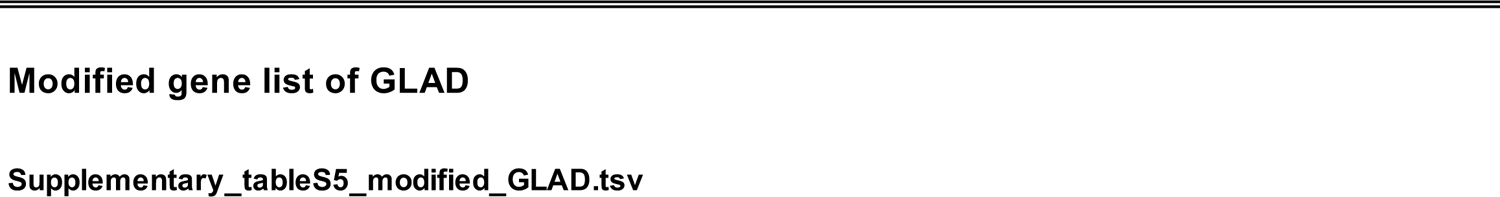

**Supplementary Table S6.**
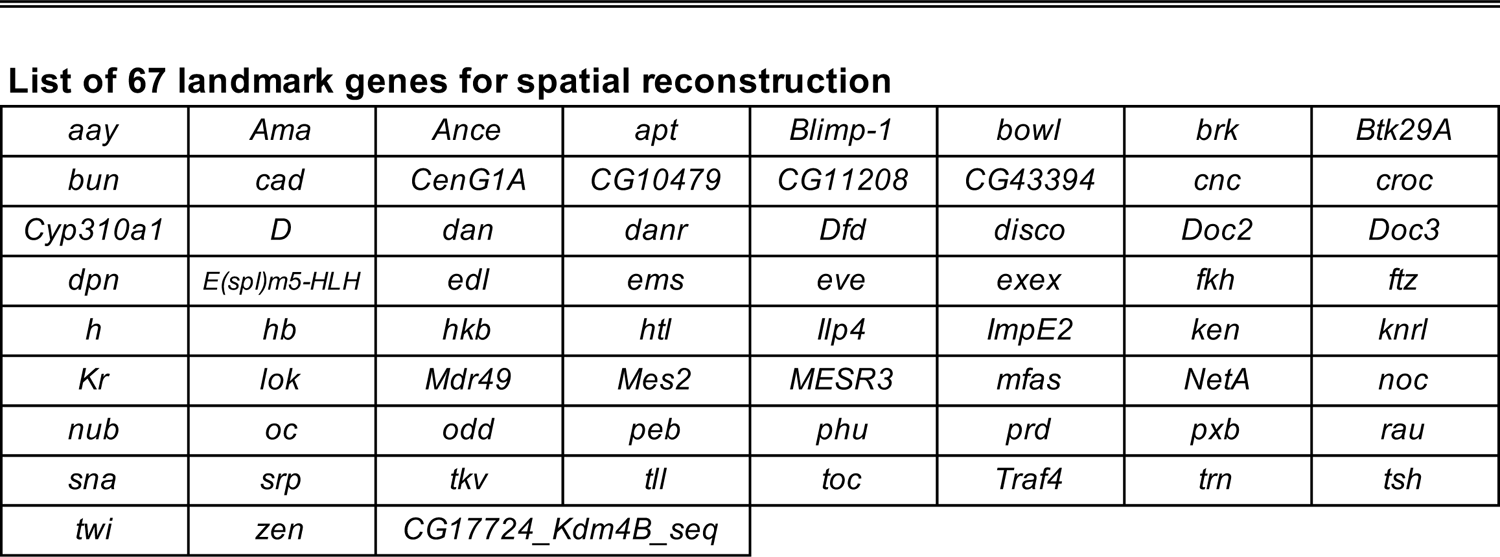

**Supplementary Figure S9:**
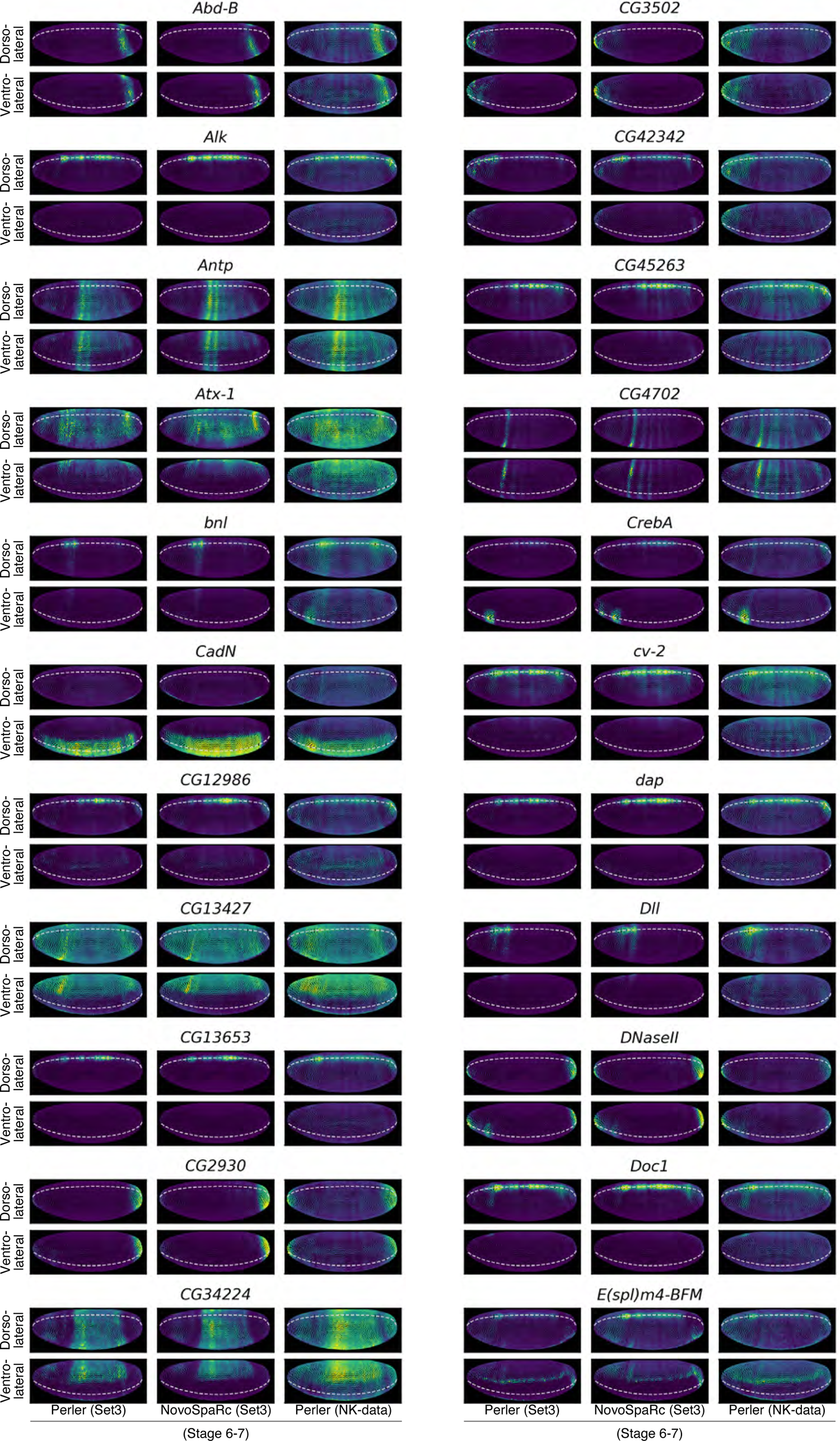
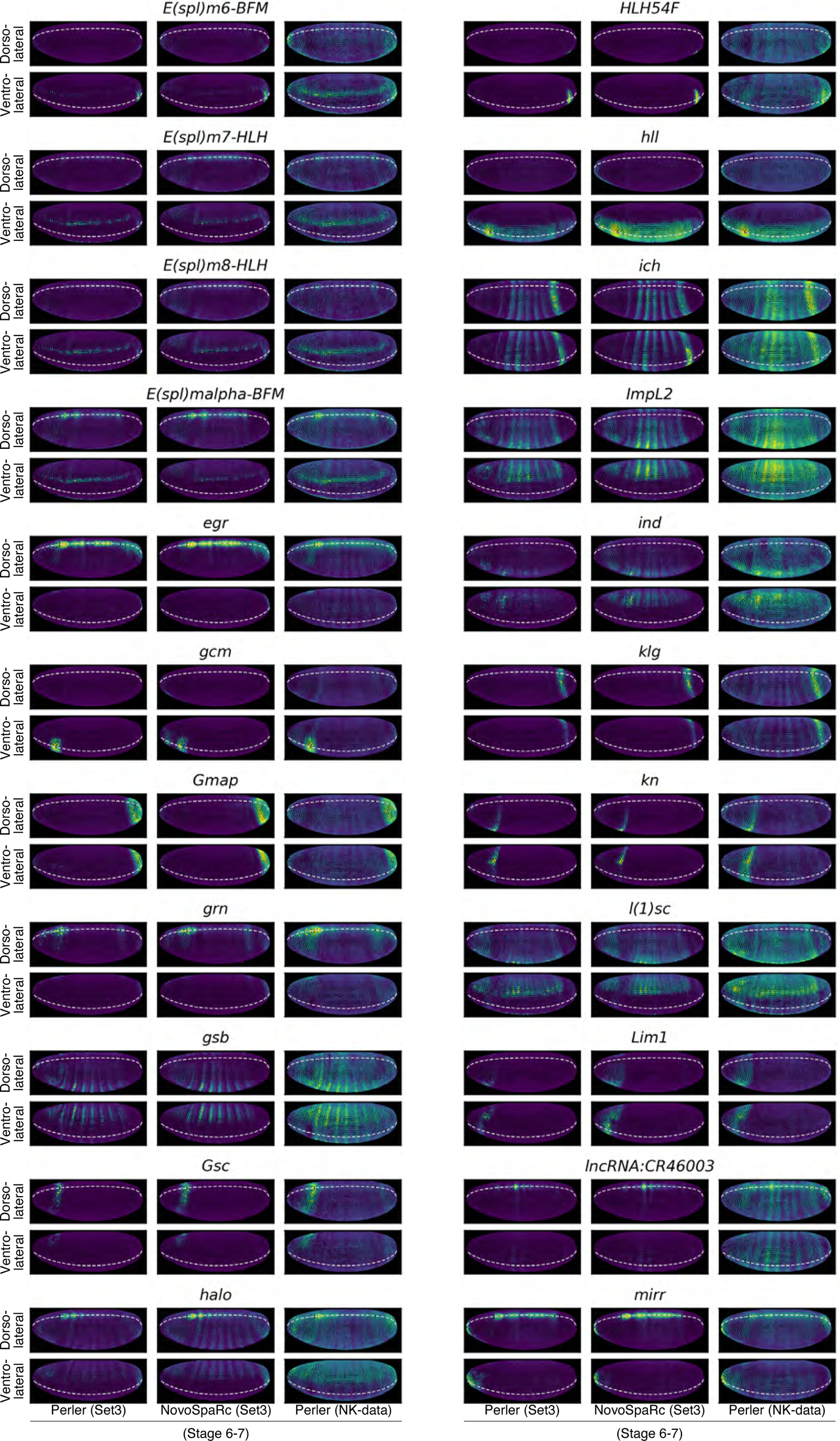
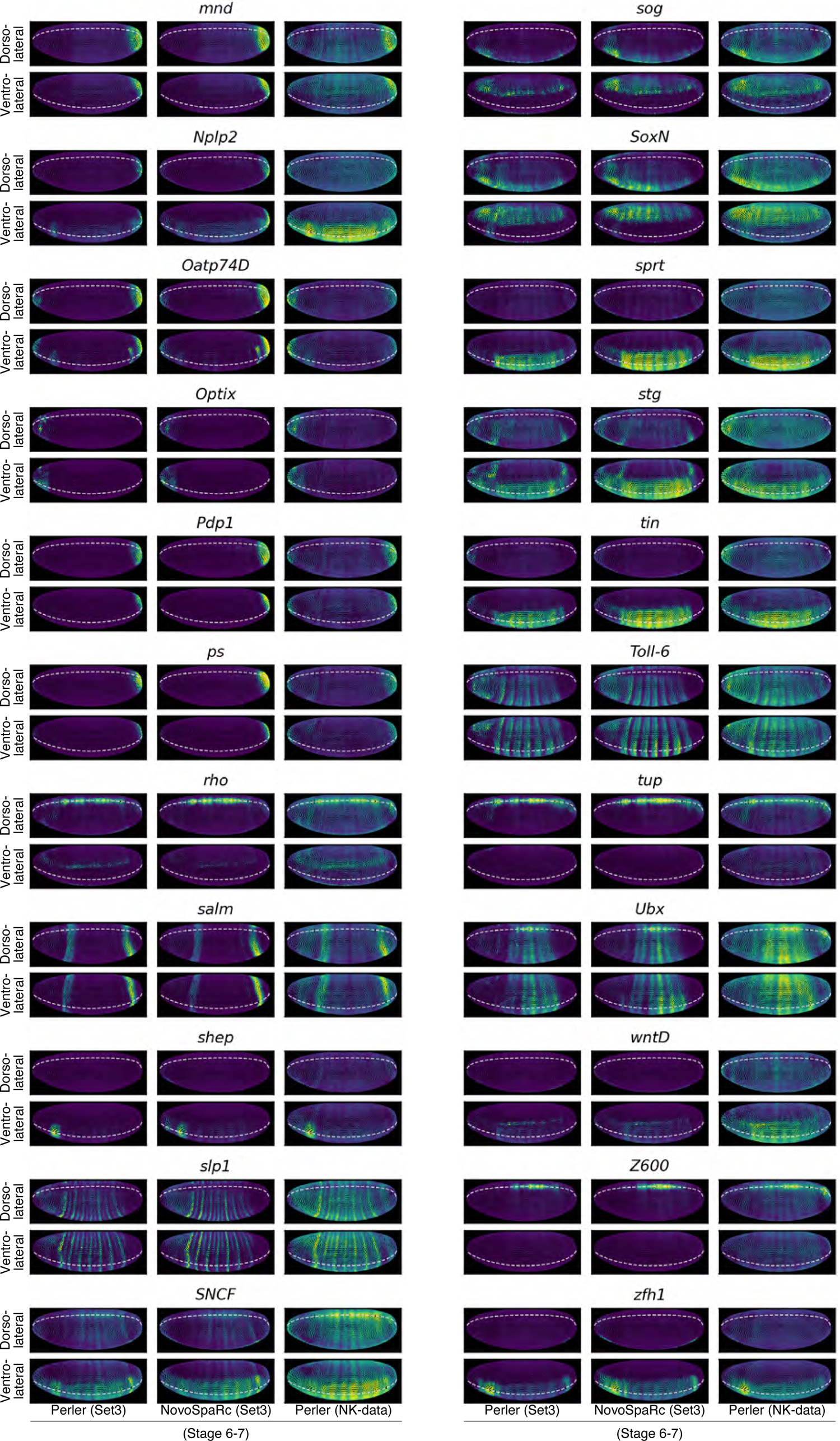
Predicted spatial patterns of non-landmark HVGs. Predictive spatial reconstruction of non-landmark 66 HVGs by Perler with Set3 data (left), NovoSpaRc with Set3 data (middle), and Perler with NK-data (right). Colormaps are linear and min-max scaled. The upper panels show the dorsolateral views (θ = 45°, Figure S7G), and the lower panels show ventrolateral views (θ = –45°, Figure S7G). Dashed lines indicate the dorsal or ventral midline. HVGs that were detected in both Set3 and NK-data and had the top 66 variances in Set3 data are shown.

## Notes

### Competing Interest Statement

The authors have declared no competing interest.

## References

1. Achim, K., Pettit, J., Saraiva, L.R., Gavriouchkina, D., Larsson, T., Arendt, D., and Marioni, J.C. (2015). High-throughput spatial mapping of single-cell RNA-seq data to tissue of origin. Nat. Biotechnol. 33, 503–509.

2. Adam, M., Potter, A.S., and Potter, S.S. (2017). Psychrophilic proteases dramatically reduce single-cell RNA-seq artifacts: A molecular atlas of kidney development. Development 144, 3625–3632.

3. Akam, M. (1987). The molecular basis for metameric pattern in the Drosophila embryo. Development 101, 1–22.

4. Del Álamo, D., Rouault, H., and Schweisguth, F. (2011). Mechanism and significance of cis-inhibition in notch signalling. Curr. Biol. 21, 40–47.

5. Bertet, C., Sulak, L., and Lecuit, T. (2004). Myosin-dependent junction remodelling controls planar cell intercalation and axis elongation. Nature 429, 667–671.

6. Bray, S., and Furriols, M. (2001). Notch pathway: Making sense of suppressor of hairless. Curr. Biol. 11, 217–221.

7. Briggs, J.A., Weinreb, C., Wagner, D.E., Megason, S., Peshkin, L., Kirschner, M.W., and Klein, A.M. (2018). The dynamics of gene expression in vertebrate embryogenesis at single-cell resolution. Science 360, eaar5780.

8. Van Den Brink, S.C., Sage, F., Vértesy, Á., Spanjaard, B., Peterson-Maduro, J., Baron, C.S., Robin, C., and Van Oudenaarden, A. (2017). Single-cell sequencing reveals dissociation-induced gene expression in tissue subpopulations. Nat. Methods 14, 935–936.

9. Briscoe, J., and Small, S. (2015). Morphogen rules: Design principles of gradient-mediated embryo patterning. Development 142, 3996–4009.

10. Cammarota, C., Finegan, T.M., Wilson, T.J., Yang, S., and Bergstralh, D.T. (2020). An Axon-Pathfinding Mechanism Preserves Epithelial Tissue Integrity. Curr. Biol. 30, 5049–5057.e3.

11. Cang, Z., and Nie, Q. (2020). Inferring spatial and signaling relationships between cells from single cell transcriptomic data. Nat. Commun. 11, 2084.

12. Chen, S., Zhou, Y., Chen, Y., and Gu, J. (2018). Fastp: An ultra-fast all-in-one FASTQ preprocessor. Bioinformatics 34, i884–i890.

13. Clark, E., and Akam, M. (2016). Odd-paired controls frequency doubling in Drosophila segmentation by altering the pair-rule gene regulatory network. Elife 5, e18215.

14. Colas, J.F., Launay, J.M., Vonesch, J.L., Hickel, P., and Maroteaux, L. (1999). Serotonin synchronises convergent extension of ectoderm with morphogenetic gastrulation movements in Drosophila. Mech. Dev. 87, 77–91.

15. Collinet, C., and Lecuit, T. (2021). Programmed and self-organized flow of information during morphogenesis. Nat. Rev. Mol. Cell Biol. 22, 245–265.

16. Cowden, J., and Levine, M. (2002). The Snail repressor positions Notch signaling in the Drosophila embryo. Development 129, 1785–1793.

17. Denisenko, E., Guo, B.B., Jones, M., Hou, R., de Kock, L., Lassmann, T., Poppe, D., Clément, O., Simmons, R.K., Lister, R., et al. (2020). Systematic assessment of tissue dissociation and storage biases in single-cell and single-nucleus RNA-seq workflows. Genome Biol. 21, 130.

18. Falo-Sanjuan, J., Lammers, N.C., Garcia, H.G., and Bray, S.J. (2019). Enhancer Priming Enables Fast and Sustained Transcriptional Responses to Notch Signaling. Dev. Cell 50, 411–425.E8.

19. Farrell, J.A., Wang, Y., Riesenfeld, S.J., Shekhar, K., Regev, A., and Schier, A.F. (2018). Single-cell reconstruction of developmental trajectories during zebrafish embryogenesis. Science 360, eaar3131.

20. Finak, G., McDavid, A., Yajima, M., Deng, J., Gersuk, V., Shalek, A.K., Slichter, C.K., Miller, H.W., McElrath, M.J., Prlic, M., et al. (2015). MAST: A flexible statistical framework for assessing transcriptional changes and characterizing heterogeneity in single-cell RNA sequencing data. Genome Biol. 16, 1–13.

21. Fowlkes, C.C., Luengo Hendriks, C.L., Keränen, S.V.E., Weber, G.H., Rübel, O., Huang, M.Y., Chatoor, S., DePace, A.H., Simirenko, L., Henriquez, C., et al. (2008). A Quantitative Spatiotemporal Atlas of Gene Expression in the Drosophila Blastoderm. Cell 133, 364–374.

22. Ganguly, A., Jiang, J., and Ip, Y.T. (2005). Drosophila WntD is a target and an inhibitor of the Dorsal/Twist/Snail network in the gastrulating embryo. Development 132, 3419–3429.

23. Gheisari, E., Aakhte, M., and Müller, H.-A.J.A.J. (2020). Gastrulation in Drosophila melanogaster: Genetic control, cellular basis and biomechanics. Mech. Dev. 163, 103629.

24. Graham, P.L., Anderson, W.R., Brandt, E.A., Xiang, J., and Pick, L. (2019). Dynamic expression of Drosophila segmental cell surface-encoding genes and their pair-rule regulators. Dev. Biol. 447, 147–156.

25. Hafemeister, C., and Satija, R. (2019). Normalization and variance stabilization of single-cell RNA-seq data using regularized negative binomial regression. Genome Biol. 20, 296.

26. Hammonds, A.S., Bristow, C.A., Fisher, W.W., Weiszmann, R., Wu, S., Hartenstein, V., Kellis, M., Yu, B., Frise, E., and Celniker, S.E. (2013). Spatial expression of transcription factors in Drosophila embryonic organ development. Genome Biol. 14, R140.

27. Heimberg, G., Bhatnagar, R., El-samad, H., Thomson, M., Heimberg, G., Bhatnagar, R., El-samad, H., and Thomson, M. (2016). Low Dimensionality in Gene Expression Data Enables the Accurate Extraction of Transcriptional Programs from Shallow Sequencing. Cell Syst. 2, 239–250.

28. Heisenberg, C., and Bellaiche, Y. (2013). Forces in Tissue Morphogenesis and Patterning. Cell 153, 948–962.

29. Hinck, L. (2004). The versatile roles of “axon guidance” cues in tissue morphogenesis. Dev. Cell 7, 783–793.

30. Hu, Y., Comjean, A., Perkins, L.A., Perrimon, N., and Mohr, S.E. (2015). GLAD: an Online Database of Gene List Annotation for Drosophila. J. Genomics 3, 75–81.

31. Hunter, J.D. (2007). Matplotlib: A 2D graphics environment. Comput. Sci. Eng. 9, 90– 95.

32. Ingham, P.W. (1988). The molecular genetics of embryonic pattern formation in Drosophila. Nature 335, 25–34.

33. Islam, S., Zeisel, A., Joost, S., La Manno, G., Zajac, P., Kasper, M., Lönnerberg, P., and Linnarsson, S. (2014). Quantitative single-cell RNA-seq with unique molecular identifiers. Nat. Methods 11, 163–166.

34. Jaeger, J. (2011). The gap gene network. Cell. Mol. Life Sci. 68, 243–274.

35. Jaeger, J., Manu, and Reinitz, J. (2012). Drosophila blastoderm patterning. Curr. Opin. Genet. Dev. 22, 533–541.

36. Kaminow, B., Yunusov, D., and Dobin, A. (2021). STARsolo: accurate, fast and versatile mapping/quantification of single-cell and single-nucleus RNA-seq data. BioRxiv 2021.05.05.442755.

37. Karaiskos, N., Wahle, P., Alles, J., Boltengagen, A., Ayoub, S., Kipar, C., Kocks, C., Rajewsky, N., and Zinzen, R.P. (2017). The Drosophila embryo at single-cell transcriptome resolution. Science 358, 194–199.

38. Keleman, K., Rajagopalan, S., Cleppien, D., Teis, D., Paiha, K., Huber, L.A., Technau, G.M., and Dickson, B.J. (2002). Comm Sorts Robo to Control Axon Guidance at the Drosophila Midline. Cell 110, 415–427.

39. Keleman, K., Ribeiro, C., and Dickson, B.J. (2005). Comm function in commissural axon guidance: Cell-autonomous sorting of Robo in vivo. Nat. Neurosci. 8, 156–163.

40. Keränen, S.V.E., Fowlkes, C.C., Luengo Hendriks, C.L., Sudar, D., Knowles, D.W., Malik, J., and Biggin, M.D. (2006). Three-dimensional morphology and gene expression in the Drosophila blastoderm at cellular resolution II: dynamics. Genome Biol. 7, R124.

41. Kolodziejczyk, A.A., Kim, J.K., Svensson, V., Marioni, J.C., and Teichmann, S.A. (2015). The Technology and Biology of Single-Cell RNA Sequencing. Mol. Cell 58, 610–620.

42. Kondo, T., and Hayashi, S. (2015). Mechanisms of cell height changes that mediate epithelial invagination. Dev. Growth Differ. 57, 313–323.

43. Kondo, T., and Hayashi, S. (2019). Two-step regulation of trachealess ensures tight coupling of cell fate with morphogenesis in the Drosophila trachea. Elife 8, e45145.

44. Lammers, N.C., Galstyan, V., Reimer, A., Medin, S.A., Wiggins, C.H., and Garcia, H.G. (2020). Multimodal transcriptional control of pattern formation in embryonic development. Proc. Natl. Acad. Sci. U. S. A. 117, 836–847.

45. Lécuyer, E., Yoshida, H., Parthasarathy, N., Alm, C., Babak, T., Cerovina, T., Hughes, T.R., Tomancak, P., and Krause, H.M. (2007). Global analysis of mRNA localization reveals a prominent role in organizing cellular architecture and function. Cell 131, 174– 187.

46. Letsou, W., and Cai, L. (2016). Noncommutative Biology: Sequential Regulation of Complex Networks. PLoS Comput. Biol. 12, e1005089.

47. Li, B., and Dewey, C.N. (2011). RSEM: accurate transcript quantification from RNA-Seq data with or without a reference genome. BMC Bioinformatics 12, 323.

48. Liu, W., Morgan, K.M., and Pine, S.R. (2014). Activation of the Notch1 stem cell signaling pathway during routine cell line subculture. Front. Oncol. 4, 211.

49. Luengo Hendriks, C.L., Keränen, S.V.E., Fowlkes, C.C., Simirenko, L., Weber, G.H., DePace, A.H., Henriquez, C., Kaszuba, D.W., Hamann, B., Eisen, M.B., et al. (2006). Three-dimensional morphology and gene expression in the Drosophila blastoderm at cellular resolution I: data acquisition pipeline. Genome Biol. 7, R123.

50. Martin, A.C. (2020). The physical mechanisms of Drosophila gastrulation: Mesoderm and endoderm invagination. Genetics 214, 543–560.

51. Martin, M. (2011). Cutadapt removes adapter sequences from high-throughput sequencing reads. EMBnet.Journal 17, 10–12.

52. McGinnis, W., and Krumlauf, R. (1992). Homeobox genes and axial patterning. Cell 68, 283–302.

53. Miyawaki-Kuwakado, A., Wu, Q., Harada, A., Tomimatsu, K., Fujii, T., Maehara, K., and Ohkawa, Y. (2021). Transcriptome analysis of gene expression changes upon enzymatic dissociation in skeletal myoblasts. Genes to Cells 26, 530–540.

54. Morel, V., and Schweisguth, F. (2000). Repression by Suppressor of Hairless and activation by Notch are required to define a single row of single-minded expressing cells in the Drosophila embryo. Genes Dev. 14, 377–388.

55. Moriel, N., Senel, E., Friedman, N., Rajewsky, N., Karaiskos, N., and Nitzan, M. (2021). NovoSpaRc: flexible spatial reconstruction of single-cell gene expression with optimal transport. Nat. Protoc. 16, 4177–4200.

56. Nitzan, M., Karaiskos, N., Friedman, N., and Rajewsky, N. (2019). Gene expression cartography. Nature 576, 132–137.

57. O’Flanagan, C.H., Campbell, K.R., Zhang, A.W., Kabeer, F., Lim, J.L.P., Biele, J., Eirew, P., Lai, D., McPherson, A., Kong, E., et al. (2019). Dissociation of solid tumor tissues with cold active protease for single-cell RNA-seq minimizes conserved collagenase-associated stress responses. Genome Biol. 20, 210.

58. Okochi, Y., Sakaguchi, S., Nakae, K., Kondo, T., and Naoki, H. (2021). Model-based prediction of spatial gene expression via generative linear mapping. Nat. Commun. 12, 3731.

59. Packer, J.S., Zhu, Q., Huynh, C., Sivaramakrishnan, P., Preston, E., Dueck, H., Stefanik, D., Tan, K., Trapnell, C., Kim, J., et al. (2019). A lineage-resolved molecular atlas of C. elegans embryogenesis at single-cell resolution. Science 365, eaax1971.

60. Papadopoulos, D.K., and Tomancak, P. (2019). Gene Regulation: Analog to Digital Conversion of Transcription Factor Gradients. Curr. Biol. 29, R422–R424.

61. Paré, A.C., Vichas, A., Fincher, C.T., Mirman, Z., Farrell, D.L., Mainieri, A., and Zallen, J.A. (2014). A positional Toll receptor code directs convergent extension in Drosophila. Nature 515, 523–527.

62. Paré, A.C., Naik, P., Shi, J., Mirman, Z., Palmquist, K.H., and Zallen, J.A. (2019). An LRR Receptor-Teneurin System Directs Planar Polarity at Compartment Boundaries. Dev. Cell 51, 208–221.E6.

63. Perrimon, N., Pitsouli, C., and Shilo, B.-Z. (2012). Signaling mechanisms controlling cell fate and embryonic patterning. Cold Spring Harb. Perspect. Biol. 4, a005975.

64. Peter A. Lawrence (1992). The Making of a Fly: The Genetics of Animal Design (Oxford, UK: Wiley-Blackwell).

65. Petkova, M.D., Tkačik, G., Bialek, W., Wieschaus, E.F., and Gregor, T. (2019). Optimal decoding of information from a genetic network. Cell 176, 844–855.

66. Rahimi, N., Averbukh, I., Haskel-Ittah, M., Degani, N., Schejter, E.D., Barkai, N., and Shilo, B.-Z. (2016). A WntD-Dependent Integral Feedback Loop Attenuates Variability in Drosophila Toll Signaling. Dev. Cell 36, 401–414.

67. Raudvere, U., Kolberg, L., Kuzmin, I., Arak, T., Adler, P., Peterson, H., and Vilo, J. (2019). G:Profiler: A web server for functional enrichment analysis and conversions of gene lists (2019 update). Nucleic Acids Res. 47, W191–W198.

68. Reeves, G.T., and Stathopoulos, A. (2009). Graded Dorsal and Differential Gene Regulation in the Drosophila Embryo. Cold Spring Harb. Perspect. Biol. 1, a000836.

69. Robinson, M.D., McCarthy, D.J., and Smyth, G.K. (2009). edgeR: A Bioconductor package for differential expression analysis of digital gene expression data. Bioinformatics 26, 139–140.

70. Satija, R., Farrell, J. a, Gennert, D., Schier, A.F., and Regev, A. (2015). Spatial reconstruction of single-cell gene expression data. Nat. Biotechnol. 33, 495–502.

71. Saxena, A., Wagatsuma, A., Noro, Y., Kuji, T., Asaka-Oba, A., Watahiki, A., Gurnot, C., Fagiolini, M., Hensch, T.K., and Carninci, P. (2012). Trehalose-enhanced isolation of neuronal sub-types from adult mouse brain. Biotechniques 52, 381–385.

72. Schaerlinger, B., Launay, J.M., Vonesch, J.L., and Maroteaux, L. (2007). Gain of affinity point mutation in the serotonin receptor gene 5-HT2Dro accelerates germband extension movements during Drosophila gastrulation. Dev. Dyn. 236, 991–999.

73. Small, S., and Arnosti, D.N. (2020). Transcriptional Enhancers in Drosophila. Genetics 216, 1–26.

74. Staller, M. V., Fowlkes, C.C., Bragdon, M.D.J., Wunderlich, Z., Estrada, J., and DePace, A.H. (2015). A gene expression atlas of a bicoid-depleted Drosophila embryo reveals early canalization of cell fate. Development 142, 587–596.

75. Stedden, C.G., Menegas, W., Zajac, A.L., Williams, A.M., Cheng, S., Özkan, E., and Horne-Badovinac, S. (2019). Planar-Polarized Semaphorin-5c and Plexin A Promote the Collective Migration of Epithelial Cells in Drosophila. Curr. Biol. 29, 908–920.e6.

76. Stuart, T., Butler, A., Hoffman, P., Hafemeister, C., Papalexi, E., Mauck, W.M., Hao, Y., Stoeckius, M., Smibert, P., and Satija, R. (2019). Comprehensive Integration of Single-Cell Data. Cell 177, 1888–1902.E21.

77. Tanay, A., and Regev, A. (2017). Scaling single-cell genomics from phenomenology to mechanism. Nature 541, 331–338.

78. Tanevski, J., Nguyen, T., Truong, B., Karaiskos, N., Ahsen, M.E., Zhang, X., Shu, C., Xu, K., Liang, X., Hu, Y., et al. (2020). Gene selection for optimal prediction of cell position in tissues from single-cell transcriptomics data. Life Sci. Alliance 3, e202000867.

79. Tetley, R.J., Blanchard, G.B., Fletcher, A.G., Adams, R.J., and Sanson, B. (2016). Unipolar distributions of junctional myosin II identify cell stripe boundaries that drive cell intercalation throughout drosophila axis extension. Elife 5, e12094.

80. Tomancak, P., Beaton, A., Weiszmann, R., Kwan, E., Shu, S.Q., Lewis, S.E., Richards, S., Ashburner, M., Hartenstein, V., Celniker, S.E., et al. (2002). Systematic determination of patterns of gene expression during Drosophila embryogenesis. Genome Biol. 3, research0088.1.

81. Tomancak, P., Berman, B.P., Beaton, A., Weiszmann, R., Kwan, E., Hartenstein, V., Celniker, S.E., and Rubin, G.M. (2007). Global analysis of patterns of gene expression during Drosophila embryogenesis. Genome Biol. 8, R145.

82. Torre, E., Dueck, H., Shaffer, S., Gospocic, J., Gupte, R., Bonasio, R., Kim, J., Murray, J., and Raj, A. (2018). Rare Cell Detection by Single-Cell RNA Sequencing as Guided by Single-Molecule RNA FISH. Cell Syst. 6, 171–179.E5.

83. Vaughen, J., and Igaki, T. (2016). Slit-Robo Repulsive Signaling Extrudes Tumorigenic Cells from Epithelia. Dev. Cell 39, 683–695.

84. Virtanen, P., Gommers, R., Oliphant, T.E., Haberland, M., Reddy, T., Cournapeau, D., Burovski, E., Peterson, P., Weckesser, W., Bright, J., et al. (2020). SciPy 1.0: fundamental algorithms for scientific computing in Python. Nat. Methods 17, 261–272.

85. Wagner, D.E., Weinreb, C., Collins, Z.M., Briggs, J.A., Megason, S.G., and Klein, A.M. (2018). Single-cell mapping of gene expression landscapes and lineage in the zebrafish embryo. Science 360, 981–987.

86. Wang, C., Dickinson, L.K., and Lehmann, R. (1994). Genetics of nanos localization in Drosophila. Dev. Dyn. 199, 103–115.

87. Welch, J.D., Kozareva, V., Ferreira, A., Vanderburg, C., Martin, C., and Macosko, E.Z. (2019). Single-Cell Multi-omic Integration Compares and Contrasts Features of Brain Cell Identity. Cell 177, 1873–1887.E17.

88. Wickham, H. (2009). ggplot2: Elegant Graphics for Data Analysis (Springer-Verlag New York).

89. Wieschaus, E., and Nüsslein-Volhard, C. (2016). The Heidelberg Screen for Pattern Mutants of Drosophila: A Personal Account. Annu. Rev. Cell Dev. Biol. 32, 1–46.

90. Wilk, R., Hu, J., Blotsky, D., and Krause, H.M. (2016). Diverse and pervasive subcellular distributions for both coding and long noncoding RNAs. Genes Dev. 30, 594–609.

91. Yazdani, U., and Terman, J.R. (2006). The semaphorins. Genome Biol. 7, 211.

92. Yoo, S.K., Pascoe, H.G., Pereira, T., Kondo, S., Jacinto, A., Zhang, X., and Hariharan, I.K. (2016). Plexins function in epithelial repair in both Drosophila and zebrafish. Nat. Commun. 7, 12282.

93. Zallen, J. a, and Blankenship, J.T. (2008). Multicellular dynamics during epithelial elongation. Semin. Cell Dev. Biol. 19, 263–270.

94. Zallen, J. a, and Wieschaus, E. (2004). Patterned gene expression directs bipolar planar polarity in Drosophila. Dev. Cell 6, 343–355.

95. Zhang, M.J., Ntranos, V., and Tse, D. (2020). Determining sequencing depth in a single-cell RNA-seq experiment. Nat. Commun. 11, 774.

96. Zinzen, R.P., Cande, J., Ronshaugen, M., Papatsenko, D., and Levine, M. (2006). Evolution of the Ventral Midline in Insect Embryos. Dev. Cell 11, 895–902.

97. Zinzen, R.P., Girardot, C., Gagneur, J., Braun, M., and Furlong, E.E.M. (2009). Combinatorial binding predicts spatio-temporal cis-regulatory activity. Nature 462, 65– 70.

